# DTK-Dengue: A new agent-based model of dengue virus transmission dynamics

**DOI:** 10.1101/376533

**Authors:** K.J. Soda, S.M. Moore, G. España, J. Bloedow, B. Raybaud, B. Althouse, M.A. Johansson, E. Wenger, P. Welkhoff, T.A. Perkins, T.A. Perkins, Q.A. ten Bosch

## Abstract

Dengue virus (DENV) is a pathogen spread by *Aedes* mosquitoes that has a considerable impact on global health. Agent-based models can be used to explicitly represent factors that are difficult to measure empirically, by focusing on specific aspects of DENV transmission dynamics that influence spread in a particular location. We present a new agent-based model for DENV dynamics, DTK-Dengue, that can be readily applied to new locations and to a diverse set of goals. It extends the vector-borne disease module in the Institute for Disease Modelling’s Epidemiological Modeling Disease Transmission Kernel (EMOD-DTK) to model DENV dynamics. There are three key modifications present in DTK-Dengue: 1) modifications to how climatic variables influence vector development for *Aedes* mosquitoes, 2) updates to adult vector behavior to make them more similar to *Aedes,* and 3) the inclusion of four DENV serotypes, including their effects on human immunity and symptoms. We demonstrate DTK-Dengue’s capabilities by fitting the model to four interrelated datasets: total and serotype-specific dengue incidences between January 2007 and December 2008 from San Juan, Puerto Rico; the age distribution of reported dengue cases in Puerto Rico during 2007; and the number of adult female *Ae. aegypti* trapped in two neighborhoods of San Juan between November 2007 and December 2008. The model replicated broad patterns in the reference data, including a correlation between vector population dynamics and rainfall, appropriate seasonality in the reported incidence, greater circulation of DENV-3 than any other serotype, and an inverse relationship between age and the proportion of cases associated with each age group over 20 years old. This exercise demonstrates the potential for DTK-Dengue to assimilate multiple types of epidemiologic data into a realistic portrayal of DENV transmission dynamics. Due to the open availability of the DTK-Dengue software and the availability of numerous other modules for modeling disease transmission and control from EMOD-DTK, this new model has potential for a diverse range of future applications in a wide variety of settings.

## INTRODUCTION

With as many as 390 million infections in a given year and no evidence of decline, dengue virus (DENV) poses an ever-increasing public health problem (Bhatt et al., 2013). Although some DENV infections remain asymptomatic or cause clinically inapparent symptoms, others come with more burdensome manifestations, including dengue fever (DF), dengue hemorrhagic fever (DHF), and dengue shock syndrome (DSS). DENV is a mosquito-borne flavivirus with four distinct serotypes. Infection with one serotype is believed to cause lifelong homologous immunity, as well as temporary heterologous cross-immunity (Sabin 1950; Sabin 1952; Reich et al. 2013). Secondary infections carry an elevated risk of severe manifestations, likely due to nonneutralizing antibodies from previous infections (Katzelnick et al. 2017). Although mild and inapparent infections are associated with lower levels of viremia, recent studies have found that these individuals too can contribute substantially to DENV transmission (Duong et al. 2015, ten Bosch et al. 2018).

*Aedes aegypti* is the primary vector of DENV. It is a diurnal, container-breeding mosquito found in urban, densely populated areas in the tropics. The eggs of *Ae. aegypti* are desiccation-resistant and only require a small amount of water to hatch and develop (Christophers 1960). As a result, they are commonly found in human-made containers such as waste materials, flowerpots, open sewers, and cisterns (Christophers 1960). The availability of these larval habitats is a major driver of the mosquito’s population dynamics and has been found to be linked to climatic factors such as rainfall and temperature (Moore *et al.* 1978, Barrera *et al.* 2011), as well as social factors such as access to piped water, quality of waste removal programs, and public awareness (Barrera *et al.* 2011). These relationships are complex and context specific. Beyond reports of positive correlations between *Ae. aegypti* population sizes and rainfall, droughts have been associated with increased population sizes due to increases in water storage (Beebe et al. 2009). Moreover, dominant breeding sites vary greatly across settings (Scott *et al.* 2000).

Efforts to control the spread of DENV would benefit greatly from high-quality, multilevel data on factors underlying its propagation. Unfortunately, many relevant local factors, such as population immunity, the proportion of unobserved infections, and the population dynamics of *Ae. aegypti,* are challenging to quantify. Mathematical models for DENV dynamics can help provide insights about these factors and explore how they interact to generate observed dynamics (Perkins *et al.* 2014). Such models need to be broad enough in scope to address the diversity of relevant factors and their interactions. Agent-based models are a class of models particularly suitable for describing systems such as DENV transmission, where dynamics emerge from the interplay of processes that span different temporal and spatial scales.

There has been some effort to model *Ae. aegypti* and DENV dynamics in an agent-based context, each model differing in its emphasis and details. On one end of a spectrum are models that focus entirely on *Ae. aegypti* life history and population dynamics. For example, Skeeter Buster tracks interconnected subpopulations of *Ae. aegypti* through egg, larval, pupal, and adult life stages in relation to larval habitat availability and its relation to climatic factors (Magori et al., 2009). Model of Mosquito Aedes (MOMA) also traces subpopulations through each life stage, but particularly focuses on more accurate portrayals of vector behavior, especially as it pertains to blood feeding and oviposition (Maneerat & Daudé, 2016). On the other end of the spectrum are models that emphasize human populations and viral transmission without explicitly modeling the mosquitoes’ pre-adult life stages. Common features of these models include human mobility, viral incubation periods, serotype-specific and heterologous immunity, differentiation between apparent and inapparent infections, and rainfall-mediated mosquito population dynamics (e.g., Chao *et al.* 2012; Hladish *et al.* 2016). Some models additionally incorporate spatial heterogeneity in mosquito populations and human attractiveness to mosquitoes (e.g., Perkins *et al.* 2016). Finally, some models strike a balance between these two extremes. For example, Karl *et al.* (2014) presented a model that was capable of including both pre-adult life stages and detailed human populations by focusing on a single serotype of DENV.

The current standard in DENV modelling is toward specialized and focused models. This means a close association between the model and a specific purpose (e.g., tracing mosquito genotypes in Magori *et al.* 2009) and/or a specific location (e.g., Cairns, Australia in Karl *et al.* 2014). Implicit to this specialization toward a certain locality is a specialization toward a specific scale as well, be it city (e.g., Magori *et al.* 2009; italic> Chao *et al*. 2012; Karl *et al.* 2014; Maneerat and Daudé 2016; Perkins *et al.* 2016), province/state (e.g., Karl *et al.* (2014)), or international (Zhang *et al.* 2017). In addition to these specialized models, though, it would be helpful to have a generalized model; i.e., one that can easily transition across locations, scales, and purposes.

Here we introduce a new climate-driven, agent-based model for DENV transmission that is flexible enough to represent a diverse set of locations at a user-selected scale but is also detailed enough to address a broad range of research questions. We call this model DTK-Dengue. DTK-Dengue belongs to a preexisting suite of models from the Institute for Disease Modeling called Epidemiological Modeling Disease Transmission Kernel (EMOD-DTK). It was created and validated by a development team that has maintained a consistent and high standard for the underlying software. EMOD-DTK has the ability to address various spatial scales, from small communities to cities and entire countries. Although EMOD-DTK already has capacities to simulate vector population dynamics and diseases, DTK-Dengue differs from existing EMOD-DTK modules through its focus on the life history of *Ae. aegypti* vectors, including egg, larval, immature, and adult life stages, and on the complexities that DENV’s four serotypes impose on immunity, infectiousness, and symptom onset. We provide an example of a DTK-Dengue implementation based on data from the San Juan-Carolina-Caguas Metropolitan Statistical Area (MSA) of Puerto Rico between January 2007 and December 2008.

## METHODS

We developed a new climate-driven agent-based model for *Aedes aegypti* vectored DENV-transmission. Using data on local climatic and demographic variables, we parameterized the model to be representative of the San Juan MSA. The model was fitted to entomological and epidemiological data using a modified steepest ascent algorithm. To increase our understanding of how processes related to viral transmission and population immunity interact to generate observed DENV dynamics, we used data from different levels of the transmission system, including annual disease incidence, the age-distribution of cases, the relative serotype abundance, and the population dynamics of female *Ae. aegypti.*

### Model description

Epidemiological Modeling software (EMOD-DTK) is an agent-based model (ABM) of pathogen transmission and infectious disease occurrence. The software tracks the states and interactions of computational agents (specifically individual humans and mosquitoes for mosquito-borne diseases) through a set of probabilistic rules. These rules determine what events (demographic or epidemiological) the individual experiences at a given time and how these events affect the individual’s state. The model is stochastic in nature, as many of the agents’ rules and behaviors are contingent upon random draws from predefined distributions. As a result, no single run is the same, and suites of simulations can be performed to sample from a range of plausible outcomes under a given scenario. The model can simulate numerous infectious diseases, including malaria (Eckhof 2011, 2012a,2012b,2013), tuberculosis (Huynh *et al.* 2015), polio (Wagner *et al.* 2014), and HIV-AIDS (Bershteyn *et al.* 2013). We adapted it to simulate the life history of *Ae. aegypti* mosquitoes and the dynamics of multi-serotype transmission of DENV infection and disease.

#### Aquatic habitat

Mosquito populations are tightly linked to the availability of aquatic habitats where adult females oviposit and eggs proceed through aquatic life stages to emerge as adults. EMOD-DTK contains representations of multiple types of aquatic habitat, each of which contribute to the overall carrying capacity (*K*) of the aquatic habitat at a given time as a function of climatic variables and/or human density (Eckhoff 2011). For *Ae. aegypti,* we incorporated two habitat types, a temporary, rain-filled habitat and a constant habitat. In the temporary, rain-filled habitat, the larval carrying capacity (*K_temp_*) per unit area (*D_cell_*) fluctuated in response to rainfall (*P_rain_*), temperature (*T_K_*), and relative humidity (*RH*)

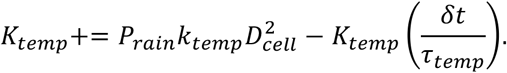

Here, the evaporation rate τ_temp_ was affected by T_A_ and RH

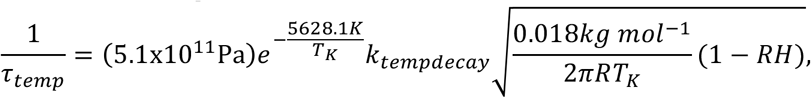

where the *k_tempdecay_* was a constant that scaled the evaporation per unit area to habitat loss. The habitat available for the simulated mosquito population was scaled by k_temp_ (further detailed under Model Fitting). Under the constant habitat, the larval carrying capacity remains fixed, regardless of weather. In combination, the constant and temporary rainfall habitats dictated that there is a baseline carrying capacity for larval *Ae. aegypti* but that this carrying capacity can be augmented based on recent climatic events.

#### Aquatic mosquito life stages

For computational efficiency, the aquatic life stages of the mosquitoes (eggs, larvae, pupae) in EMOD-DTK were not modeled on an individual basis but rather as cohorts that advanced towards the adult stage. Each aquatic stage of *Ae. aegypti* had base development and mortality rates that could be further adjusted based on habitat availability and temperature. The base daily mortality rate for eggs was 1% and was unaffected by rainfall or temperature (Romeo Aznar *et al*. 2013). Eggs hatched at a daily Arrhenius temperature-dependent rate 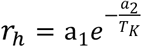 (fitted to laboratory data presented in Focks *et al.* 1993) (SFig. 1A). The model had the ability to reduce *r_h_* by a drought reduction factor of 0.33 if available habitat dropped to zero, either due to persistent drought or through control programs focused on habitat destruction (Otero, Schweigmann, and Solari 2008); because aquatic habitat had a constant component, though, this feature was never invoked. When the larval population exceeded the carrying capacity (*K*), *r_h_* was reduced by a factor relative to the larval count (*L*) and the number of eggs in the cohort ready to hatch (*δE*) to (*K — L*)/*δE*, but never by more than the drought reduction factor. Upon hatching, the base daily mortality rate for the larvae was 12%, but was further adjusted to account for density-dependence, such as would arise through cannibalism of 1st instar larvae by 4th instars or limited food availability. Whenever the daily survival resulted in the larval population exceeding K, larval mortality was increased by the degree of overpopulation *L* * *K*~^1^. Surviving larvae emerged via a pupal stage (not explicitly modeled) into adult mosquitoes at a third Arrhenius temperature-dependent rate *r_e_* (fitted to laboratory data from Yang *et al.* 2009) (SFig. 1B).

**SFig 1:**
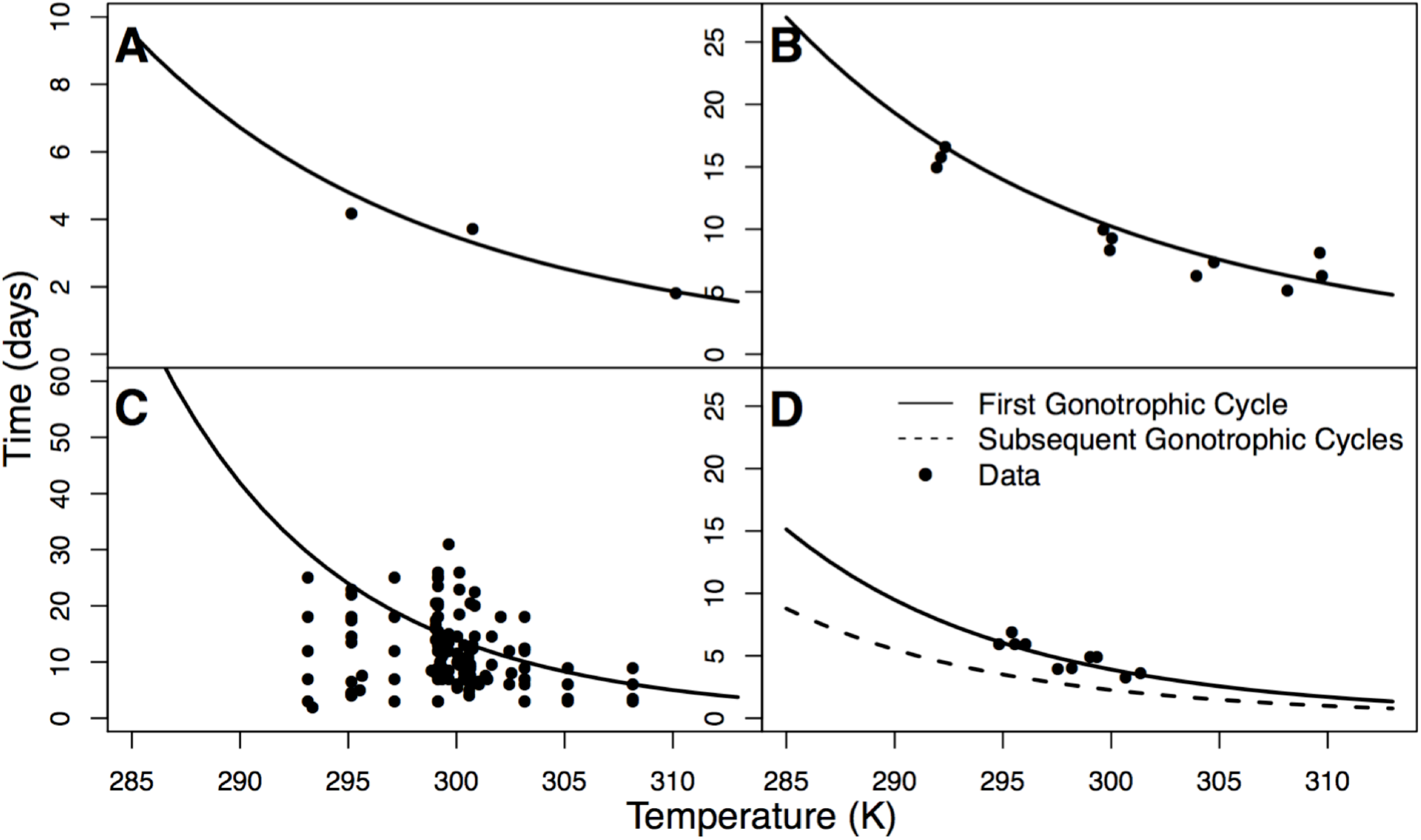
Functional forms for mosquito life traits relative to reference data from laboratory studies (black dots) with A) duration of egg hatching, B) duration of development from aquatic stages to adult, C) extrinsic incubation period, and D) gonotrophic cycle length for the first cycle (solid line) and subsequent cycles (dashed line).

#### Adult mosquito populations

Upon emergence, adult mosquitoes matured for a fixed duration of two days before they took their first blood meal. Since only female mosquitoes were modeled, adults subsequently entered a cycle of host-seeking, feeding, and egg-laying similar to the *Anopheles* implementations (Eckhoff 2011). The average duration of the first gonotrophic cycle (GC) was modeled as the inverse of a temperature-dependent Arrhenius rate *r_GC_* (fitted to data from MacDonald 1956; McClelland 1971; Pant and Yasuno 1973; Nayar 1981) (**Error! Reference source not found.D**). After the first cycle, this length was reduced by 58% (Focks *et al.* 1993). During each cycle all surviving adult mosquitoes experienced an event, the nature of which was decided via a decision tree (see Eckhoff (2011) for complete tree). Outcomes were categorizable into no action, ovipositioning, or death. To oviposition, the mosquito must find a host and take a blood meal. While a mosquito could feed on a non-human host, *Aedes aegypti* fed predominantly on humans (anthropophilic fraction: 95%) (Scott *et al.* 1993). Further, *Ae. aegypti* were assumed to feed indoors (Scott *et al.* 2000) and during the day (Chadee and Martinez 2000), decisions that could affect the effect of intervention programs (not used in this study). Upon biting, a mosquito experienced an elevated hazard of dying (10%) (Day *et al.* 1994). In the event of a successful bite on an infectious individual, a susceptible mosquito acquired a DENV infection with a probability determined by the time-varying infectiousness of humans (SFig 2) (ten Bosch *et al.* 2018, *submitted).* Infected mosquitoes experienced an extrinsic incubation period (EIP) whose duration depended on a temperature-dependent Arrhenius function. Initial parameters for this Arrhenius function were fit to data from Chan and Johansson (2012) (**Error! Reference source not found.C**) but were further adjusted based on epidemiological data (see Model Fitting). After the EIP, an infected mosquito became infectious and could spread DENV through bites on susceptible humans at a predefined probability. Following a bite, the mosquito entered a resting period after which it started looking for a hatching site to lay eggs. The adult lifespan was exponentially distributed with an average of 14 days (Christophers 1960) but was additionally affected by interventions and hazards experienced during the feeding cycle.

**SFig 2:**
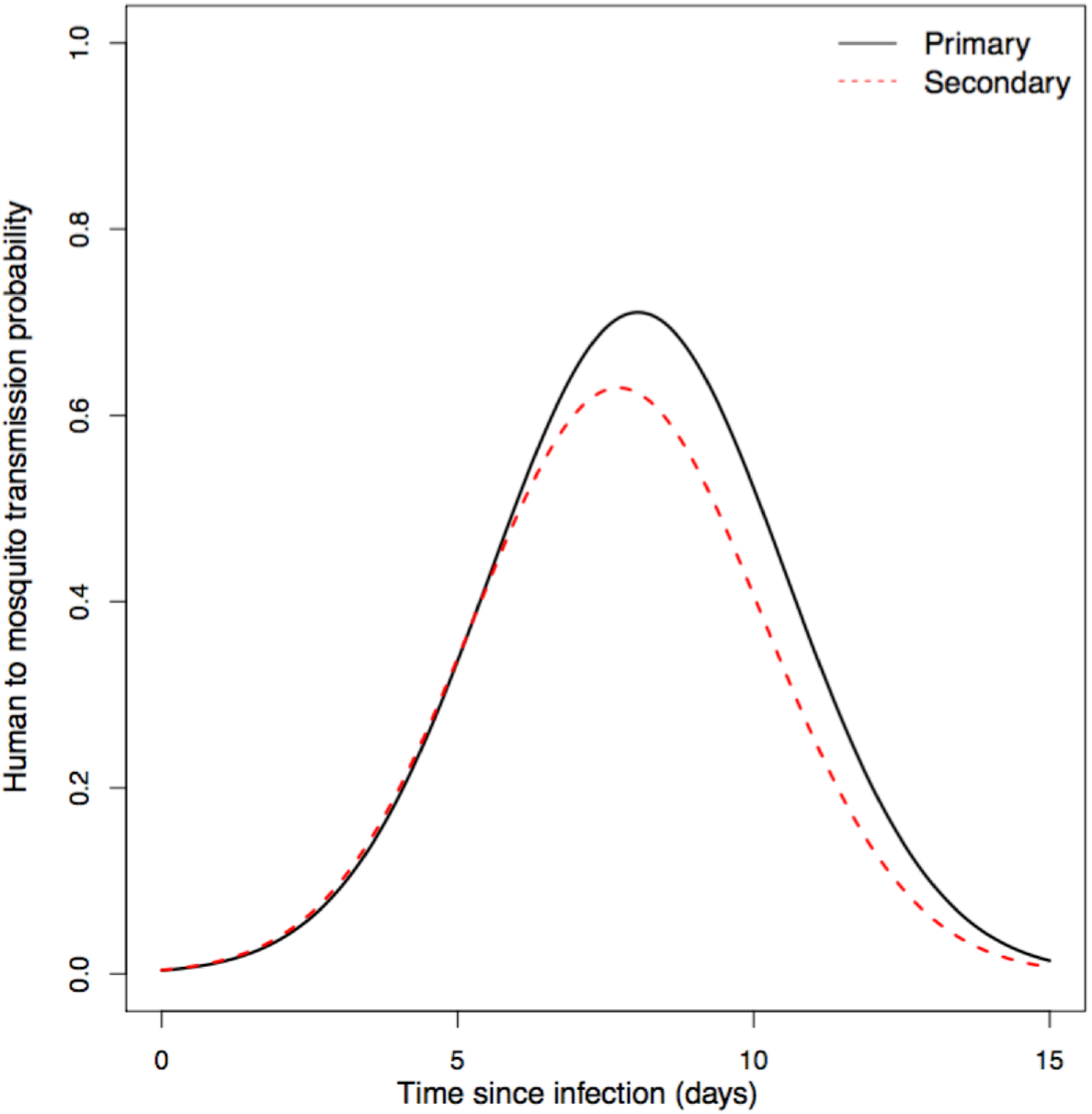
Human infectiousness to mosquitoes over time

#### Human population

Each human had a set of traits representing its demographics (age, sex, and the relative risk of being bitten), and serotype-specific health status (infection status, immunity, presence/absence of symptoms, and health-seeking behavior). At the onset of the simulation, the pre-existing serotype-specific immunity of each human was informed by a random draw from an exponential distribution with mean 1 – exp (*Λ_i_* * *age*), where *Λ_i_* is the time-averaged annual force of infection each human experienced over the course of its lifetime. Throughout the simulation, humans aged and died based on age- and sex-dependent mortality rates (see Data). New, immunologically naive individuals were also born into the population based on the current population size and the crude birth rate. Simulations were tailored to specific localities through empirical estimates of the mortality and crude birth rates (see Data), as well as the initial age distribution of the population.

#### Human infection

Each human was assumed to have an equal probability of encountering each mosquito in its population (i.e., homogeneous mixing). The risk of being bitten depended on age, with the biting risk increasing up to the age of twenty and remaining stable thereafter (Liebman *et al.* 2014). To account for heterogeneity within ages, each human received an exponentially distributed scaling factor (mean: 1) that further adjusted the individual’s biting risk. Upon infection, the probability that an infected human progressed to symptomatic disease depended on the number of previous infections the individual experienced (primary infection: 18%; secondary infection: 24%; postsecondary infection: 9.2%) (Clapham 2017; ten Bosch *et al.* 2018, *submitted).* Symptomatic infections then had an 8% probability of seeking health care and consequently being reported (Stanaway et al. 2016). In these cases, there was a lognormally distributed delay between infection and reporting (log mean: 2, log width: 0.27) (Tomashek *et al.* 2012). After recovery, humans experienced an exponentially distributed period of cross-immunity to all serotypes. Thereafter, the individuals remained immune solely to those serotypes to which they had been exposed.

The probability that a susceptible mosquito became infected upon biting an infected person changed over the course of a person’s infection, with primary infections being somewhat more infectious than post-primary infections (SFig. 2). These differences reflect a recent synthesis (ten Bosch et al. 2018, *submitted)* of within-host viremia models and viremia-infectiousness relationships.

Spatial structuring of human and mosquito populations is represented in EMOD-DTK by sub-dividing the population into separate nodes. Each spatial node in the model can have unique climate values, larval mosquito habitat amounts, and human population sizes. The mosquito population dynamics and transmission processes described above occur within a node, with nodes connected by human and mosquito movement. The spatial scales considered in this study are much larger than the typical dispersal distance of *Ae. aegypti* (Harrington et al. 2005), so the possibility of internode movement by mosquitoes was ignored. Parameter values were the same for each spatial node, except the estimated amount of larval mosquito habitat was scaled by the human population size in each node.

### Data

#### Study Area and Climate data

The San Juan-Carolina-Caguas Metropolitan Statistical Area (MSA) is a roughly 3,983 km^2^ region in northeastern Puerto Rico that is composed of 40 municipalities. The population in 2007 was estimated to be 2,382,377. Each municipality in the San Juan MSA was represented as a separate node within a multi-node simulation. The relative population size of each municipality reflected 2010 WorldPop estimates (available at www.worldpop.org) (Sorichetta *et al.* 2015), but, to reduce the computational cost of each simulation, each population was scaled to one quarter its actual size. Human movement between municipalities was modeling using a gravity model, with movement rates between municipalities proportional to the population size of each locality, and inversely proportional to the distance between municipalities. The gravity model parameter values were obtained from a study that fit a gravity model to county-level commuter data in the United States (Viboud et al. 2006). A proportionality constant scaling the gravity-model movement rates to between-municipality movement rates in Puerto Rico was estimated as part of the model fitting process. The movement rates were represented in the model as the fraction of individuals in municipality *i* who travelled in a given day to municipality j. Movement events were assumed to last for one day, with a 100% probability that the individual would return to their home municipality.

Climate maps of air temperature, rainfall, and relative humidity were created for 2006–2008 using the algorithm in Chabot-Couture, Nigmatulina, and Eckhoff (2014). Briefly, the algorithm uses temperature anomaly and dew point data from the Global Summary of the Day database (Global Summary of the Day 2012) and rainfall data from NOAA STAR’s Hydro-Estimator (Scofield and Kuligowski 2003) to interpolate these climatic variables to nearby locations at a 2.5-arcminute resolution through a combination of Kriging and bilinear interpolation. The temperature anomalies were then converted to air temperatures based on climate norms from WorldClim (Hijmans *et al.* 2005), and the dew point data was converted to relative humidity based on region-specific lapse rates and chemophysical equations. To generate a single set of climatic variables for each municipality in the San Juan MSA, the climate maps were segmented into individual municipalities, and daily, population-weighted means of each climatic variable were taken.

#### Epidemiological and entomological data

The Puerto Rico Department of Health and the Centers for Disease Control and Prevention provided weekly DENV-case incidences for the San Juan MSA between January 1^st^, 2007 and December 31^st^, 2007, and these data were then aggregated to month (Fig. 1). A total of 1,969 DENV cases were reported by the Passive Dengue Surveillance System (PDSS) over this time frame. Cases were generally laboratory confirmed, except when laboratory capacity was exceeded during high transmission periods. Confirmation was done using RT-PCR or MAC-ELISA, depending on the timing of the blood sample relative to symptom onset. The serotype was identified for 61.6% of confirmed cases. The data on serotype occurrences reflected the relative (co-)dominance of the different serotypes over time (Fig. 2). All incidence data is available through the Dengue Forecasting Challenge’s website (http://dengueforecasting.noaa.gov/).

**Fig. 1:**
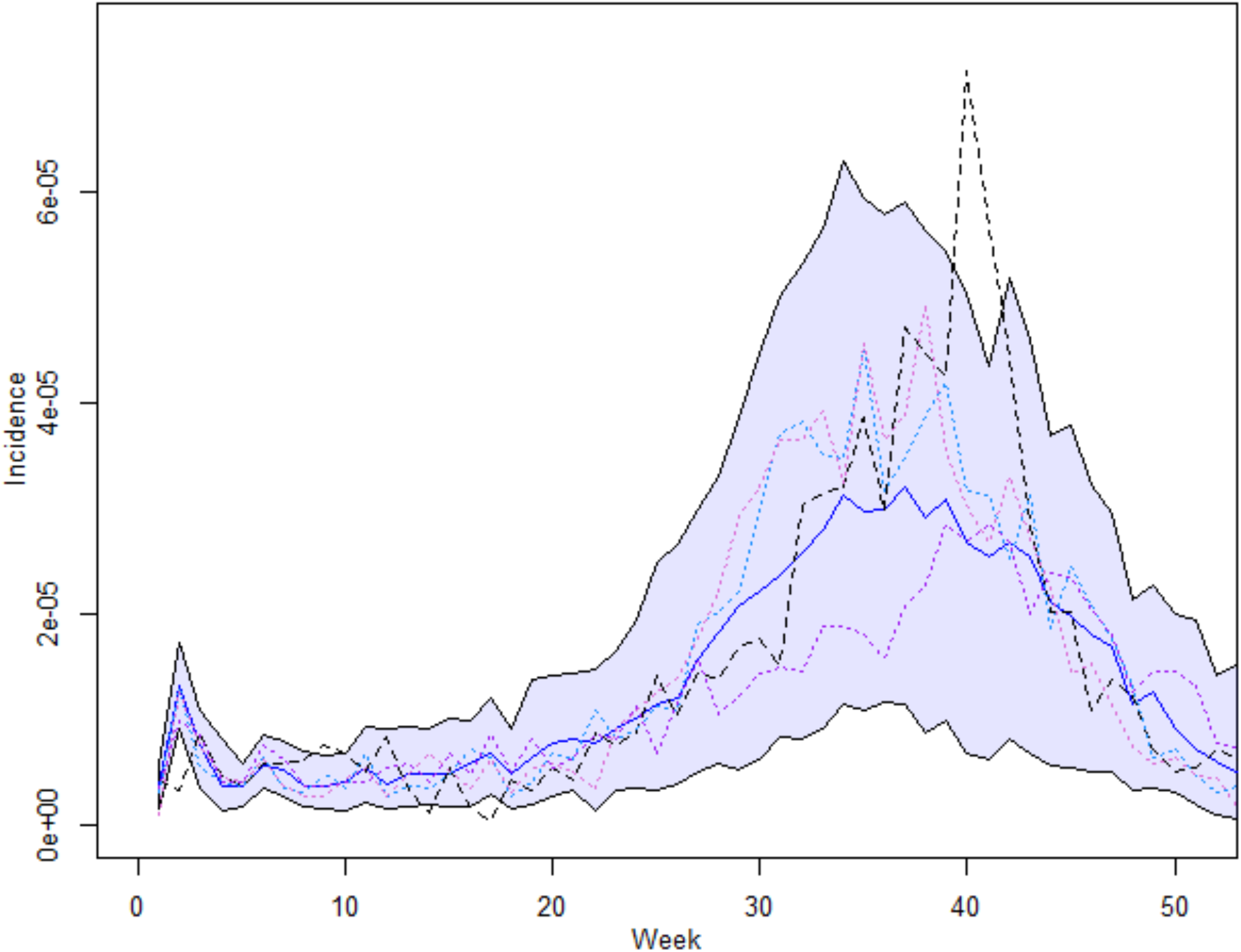
Weekly reported incidence of dengue in the San Juan Metropolitan Statistical Area between Jan 1, 2007 and Dec 31, 2007 (dashed black line), along with the median reported incidence across replicate simulations (solid blue line) and 95% confidence intervals (blue area). Colored, dotted lines are example replicate simulations.

**Fig. 2:**
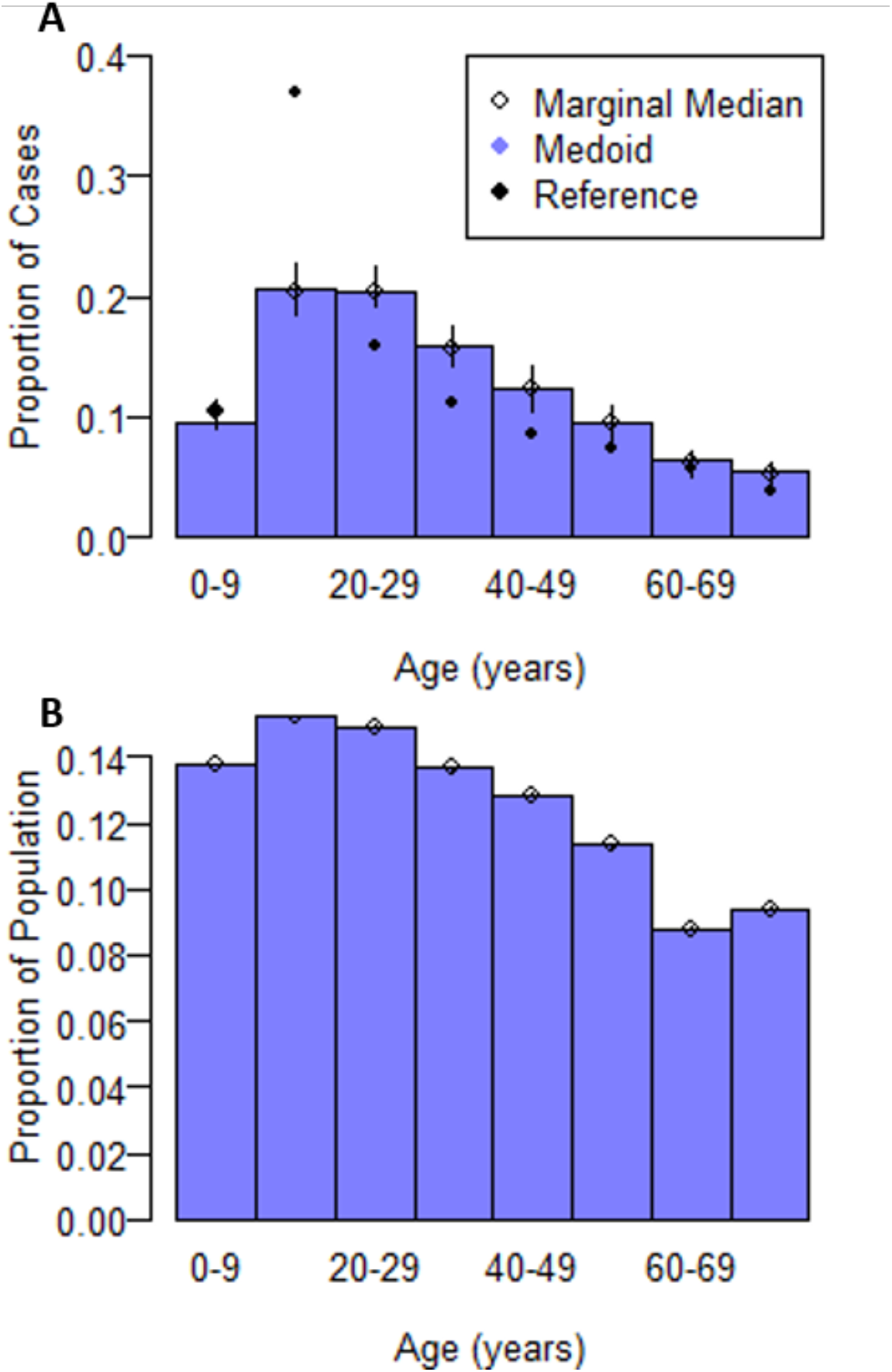
A) The age distribution of reported cases across replicate simulations of the maximum likelihood parameter set. Purple bars represent the medoid age distribution. Open circles (o) indicate the (marginal) median proportion of cases associated with each age group along with 95% confidence intervals (lines). Filled circles (•) provide the actual proportion of cases associated with each age group between January and December 2007 island-wide (Tomashek *et al.* 2001). B) The population’s age distribution across replicate simulations of the maximum likelihood parameter set. Purple bars represent the medoid age distribution. Open circles (o) indicate the marginal median proportion of the population within each age group.

Additional data on the age distribution of reported cases and the relative population sizes of female *Ae. aegypti* supplemented the PDSS data. The island-wide age distribution of laboratory-confirmed cases reported by PDSS in 2007 was derived from Tomashek *et al.* (2009), and the age bins were consolidated into ten-year blocks. We assumed that the age distribution in the San Juan MSA matched the island-wide distribution. Further, Barrera *et al.* (2011) recorded the number of female *Ae. aegypti* that were captured in BG traps within two neighborhoods of the San Juan and Carolina municipalities between November 2007 and December 2008. We assumed that *Ae. aegypti* population dynamics in each municipality reflected the abundance data from these two locations.

#### Demographic data

Each municipality was assumed to have identical sex- and age-dependent mortality rates and crude birth rates. The crude birth rate was the average rate for Puerto Rico between 1990 and 2010 according to the 2017 Revision of World Population Prospects (United Nations, Department of Economic and Social Affairs, Population Division, 2017). The mortality rates were twice the average rate between 1990 and 2010; these rates were selected to achieve more realistic population growth within the simulations. Further, the initial age distribution in each simulation emulated that of Puerto Rico in 2007 and was assumed to be homogenous across municipalities.

Due to the nature of how likelihoods were assigned to model parameters, there was no requirement that the total population of the San Juan MSA in the simulations exactly match the population of the San Juan MSA used in the reference data, but there was an assumption that the population sizes used in the reference closely corresponded with the reported incidence in the reference (see Likelihood of the model given the data). As a result, the reference population size was based on the 2007-2008 population estimates provided during the Dengue Forecasting Challenge, rather than the 2010 estimates from WorldPop. The reference population for each month was determined via a linear interpolation between the 2007 estimate (2,382,377) and the 2008 estimate (2,369,802). In practice, the Forecasting Challenge estimates were similar to the WorldPop estimate (2,306,052). All comparisons between simulation outputs and reference data were standardized or scaled to compensate for differences in population size.

### Simulation setup

We simulated the San Juan MSA on a daily scale between Jan. 1, 2007 and Dec. 31, 2007. To ensure that that the population dynamics of the mosquitoes were realistic before the simulation began the simulations also had a 365 day burn-in period. Further, to establish an infectious pool of humans and mosquitoes at the beginning of the simulation, humans were randomly infected 20 days before the end of the burn-in based on fitted probabilities. Humans in all municipalities had an equal probability of infection, although serotypes differed in their probabilities. Finally, since the mosquito trap data was predominantly associated with 2008, rather than 2007, the simulations were allowed to continue until Dec. 31, 2008; however, only mosquito population dynamics were assessed during this period. Finally, since the mosquito trap data was predominantly associated with 2008, rather than 2007, the simulations continued until Dec. 31, 2008; however, only mosquito population dynamics were assessed during this period. The outputs from the simulations were combined over all nodes, although the number of new infections (reported or otherwise) and climatic variables were recorded for each node.

### Likelihood of the model given the data

To assign log-likelihoods to sets of DTK-Dengue parameters, a set of simulations was run using those parameters, the outputs within a simulation were combined across all nodes, the combined outputs were averaged across runs, and the averages informed a collection of likelihood functions. Each of the reference data’s four components (disease incidence, serotype dominance, age-structured disease incidence, and mosquito trap data) had its own likelihood function, and the parameter set’s total likelihood was the product of these four component functions evaluated at their corresponding values in the reference data.

Each month’s incidence was assumed to follow a binomial distribution with the number of trials equal to the population of the San Juan MSA and the probability of success equal to the monthly reported incidence in the average simulation. However, due to stochasticity across simulation runs, we modeled the binomial probabilities as beta random variables such that the disease incidence’s likelihood function was based on a beta-binomial distribution. Before any simulation was run, there were no prior assumptions about the probabilities’ true values, so the underlying beta distributions were uninformative (i.e., α_t_ = 1, β_t_ = 1). A Bayesian update adjusted α and β based on the total number of humans that were alive on any day of month *t* in the average simulation, 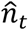, and the simulated incidence in month 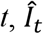, such that 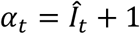 and 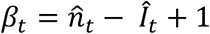 (Hobbs and Hooten 2015). Combining data across months 1 through 12, the log-likelihood based on the incidence data was:

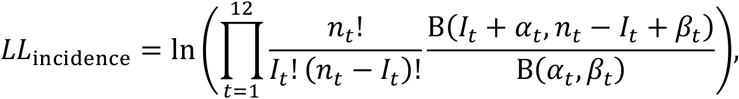

where *n_t_* and *I_t_* were the population and incidence, respectively, in month *t* of the reference data, and B(*x,y*) is the beta function.

Given the total monthly incidence and the probabilities that a case belonged to a particular serotype, we assumed that monthly serotype-specific incidence was a multinomial random variable. To incorporate between-run stochasticity, the probabilities that a case was associated with a particular serotype was assumed to be Dirichlet distributed, so the likelihood based on serotype-specific incidence was based on a Dirichlet-multinomial distribution. Similar to the disease incidence likelihood function, the Dirichlet distribution was initially uninformative, with all the hyperparameters for month *t, α_i,t_*, set to one. A Bayesian update adjusted each *α_i,t_* based on the incidence of serotype *i* during month *t* of the average simulation, 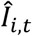, such that 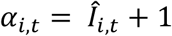 (Hobbs and Hooten 2015). Incorporating data across months 1 to 12, the log-likelihood based on the dominance data was:

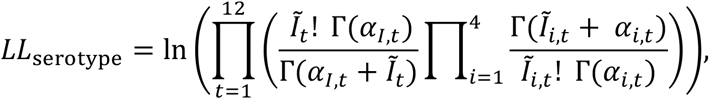

where 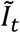 is the total number of serotyped cases in month *t* of the reference, 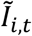 is the number of serotyped cases in month *t* of the reference that belonged to serotype *i, α_I,t_* is the sum of all *α_i,t_* for month *t*, and Γ(*x*) is the gamma function.

The likelihood function associated with the age-structured incidence data was a second Dirichlet-multinomial distribution, where the multinomially distributed variable was the number of cases associated with an age group, and the Dirichlet distributed variables were the probabilities that a case occurred in each age group. Unlike above, though, the data was aggregated across all of 2007, rather than monthly. This system was chosen to emulate how the data appeared in Tomashek *et al.* (2009). The Bayesian updates and log-likelihood calculations were otherwise comparable to that of the serotype-specific incidence. If 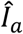 was the reported incidence in age group *a* of the average simulation, 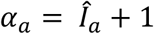 and *α_A_* is the sum of all *α*_a_. Similarly, if *I_a_*, was the incidence in age group *a* of the reference data, *I_A_* was the total incidence reported in Tomashek *et al.* (2009) data, and *A* was the total number of age bins:

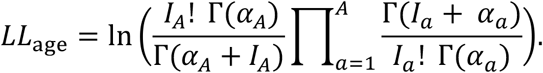

Finally, the number of trapped mosquitoes was assumed to follow a Poisson distribution. Unlike the other three likelihood components, the likelihood component for the trap data was not based on a compound distribution. Instead, the average number of adult vectors in month *t* of the average simulation, 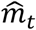, was assumed to be directly proportional to the total number of trapped mosquitoes in the San Juan and Carolina municipalities. The exact ratio between simulated and actual mosquitoes, along with the probability that a mosquito was trapped, was incorporated into a parameter, *c_trap_*, such that the expected number of trapped mosquitoes in month *t* was 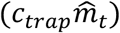. Therefore, the log-likelihood based on the trap data was:

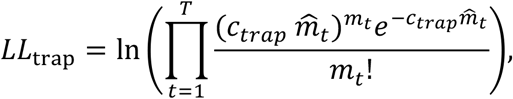

where *m_t_* is the number of trapped mosquitoes in the reference data and *T*is the total number of months for which trap data is available. Unlike the other three components, this likelihood function was evaluated for November 2007 through December 2008. After each log-likelihood component was calculated, they were summed to get a final log-likelihood for the parameters.

### Model fitting

The model was fitted to the different data sources using a maximum-likelihood approach. We fitted 16 model parameters (the carrying capacity of *Ae. aegypti* larvae in a 1×1 degree block of constant habitat, the carrying capacity of larvae in a 1×1 degree block of rain-filled habitat (*k_temp_*), the scaling constant on the evaporation rate (*k_tempdecay_*), the mosquito-to-human transmission probability, the pre-exponential factor in the extrinsic incubation period’s Arrhenius equation (i.e., *a*_1_ in 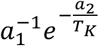), the historical force of infection for each serotype (*λ_i_*), the average duration of cross-immunity between strains, the expected number of infections of each serotype at the beginning of the simulation, the scaling parameter to convert the population size of simulated mosquitoes to the number caught in BG trap (*c_trap_*), and a scaling parameter on the migration rate) (Table 1).

**Table 1:** The boundaries for each fitted parameter, along with its maximum likelihood (ML) value.

| Parameter | Boundaries | ML Value |
| --- | --- | --- |
| Larval Carrying Capacity in Constant Habitat per Unit Area | $[1 \times 10^3, 1.5 \times 10^9]$ | $9.34 \times 10^8$ |
| Larval Carrying Capacity in Rain-Filled Habitat per Unit Area ($k_{temp}$) | $[1 \times 10^3, 5 \times 10^9]$ | $3.72 \times 10^9$ |
| Scaling Constant on Evaporation Rate ($k_{tempdecay}$) | $[0.001, 0.7]$ | 0.25 |
| Mosquito-to-Human Transmission Rate | $[0.1, 1]$ | 0.58 |
| Pre-Exponential Factor for the Extrinsic Incubation Period ($a_1$) | $[4 \times 10^{12}, 13.38 \times 10^{12}]$ | 8.04 |
| Historical Force of Infection (DENV-1) ($\lambda_1$) | $[0.001, 0.05]$ | 0.0246 |
| Historical Force of Infection (DENV-2) ($\lambda_2$) | $[0.001, 0.05]$ | 0.0355 |
| Historical Force of Infection (DENV-3)<br>($\lambda_3$) | [0.001, 0.05] | 0.0148 |
| Historical Force of Infection (DENV-4)<br>($\lambda_4$) | [0.001, 0.05] | 0.0242 |
| Mean Duration of Cross-Immunity | [180, 730] | 703.46 |
| Probability of DENV-1 Infection at Initialization | $[4 \times 10^{-6}, 1.36 \times 10^{-3}]$ | $5.04 \times 10^{-4}$ |
| Probability of DENV-2 Infection at Initialization | $[4 \times 10^{-6}, 1.36 \times 10^{-3}]$ | $9.75 \times 10^{-4}$ |
| Probability of DENV-3 Infection at Initialization | $[4 \times 10^{-6}, 1.36 \times 10^{-3}]$ | $8.89 \times 10^{-4}$ |
| Probability of DENV-4 Infection at Initialization | $[4 \times 10^{-6}, 1.36 \times 10^{-3}]$ | $7.00 \times 10^{-4}$ |
| Probability that an Adult Vector will be Caught in a BG Trap (on a log scale) | [-20, -6.9] | -10.22 |
| Scaling Parameter on the Migration Rate | [0.001, 2] | 1.07 |

The fitting algorithm was a modified steepest ascent algorithm called OptimTool, which is provided with the standard DTK-EMOD software. Within OptimTool, parameter values are constrained to be within preset bounds (Table 1). At each iteration of the algorithm, the gradient of the likelihood function was numerically estimated at a candidate solution. To do so, 100 sets of parameters were sampled around the candidate solution based on an uncorrelated multivariate normal distribution; any parameter value that exceeded its bounds were replaced with the boundary value. The log-likelihood of each parameter set was estimated based on five replicate simulations. Then, the log-likelihood of each parameter set was regressed onto the sample values. If the coefficient of determination for this regression was greater than 0.5, the regression coefficients were used as an approximation to the gradient, and the algorithm updated the candidate solution by moving in this direction in parameter space. Otherwise, the candidate solution was set to whichever parameter set had the greatest likelihood. This process was repeated for at least five iterations. If, after five iterations, the candidate solution did not remain at the same location for at least two consecutive iterations, the algorithm was restarted using a smaller mean for the multivariate normal distribution until this criterion was met, at which point the candidate was determined to be the maximum likelihood solution. The initial candidate solution was generally the midpoint of each parameter’s bounds; the sole exception was the preexponential factor in the extrinsic incubation period’s Arrhenius equation, which started at 10.

### Model performance

Since DTK-Dengue is a stochastic model, each simulation run may provide different outputs, even under the same parameter values. To explore the range of possible outcomes under the maximum likelihood solution, we ran 100 simulations under this solution. Since the likelihood function was evaluated based on the average of five simulations, we then randomly generated 100 groups of five to create 100 average simulation results. Based on these average simulations, 95% confidence intervals were placed around outputs of interest based on the values falling between the 2.5^th^ and 97.5^th^ quantiles.

## RESULTS

### Model convergence and maximum likelihood parameter values

The fitting algorithm ran for five iterations, during which the candidate solution shifted positions at every iteration. The multivariate normal distribution’s mean was then reduced by one third, and the algorithm was restarted at the parameter set with the greatest likelihood in the first five iterations. After the fourth iteration, the candidate solution remained at the same parameter values, and this parameter set was declared the maximum likelihood (ML) solution.

Table 1 reports the ML value for each fitted parameter. The values of those parameters which have direct interpretations are generally plausible. For example, the habitat decay rate parameter was estimated to be 0.255. This means that at the region’s average temperature (24.29 °C) and relative humidity (82.18%) in 2007, the larval carrying capacity within rain-filled habitats would decrease to half its size in around 4.2 days if no additional precipitation were to occur. Further, the larval carrying capacity per unit volume is an order of magnitude larger for rain-filled habitats than for constant habitats, consistent with the close association between *Ae. aegypti* and rain-filled containers. Finally, the mean duration of cross-immunity is similar to the 680-day estimate provided in Reich *et al.* (2013). However, even for parameters whose values are not well established, such as the mosquito-to-human transmission rate, estimated values are located away from the boundaries of their allowable ranges, indicating a tendency against extreme values. Alternative starting conditions and strategies for initializing simulations led to different local maxima, but these parameter sets led to simulations where certain output streams were biologically unrealistic, such as the extrinsic incubation period (EIP) (SFig. 3) or the population immunity (SFig. 4).

**SFig. 3.**
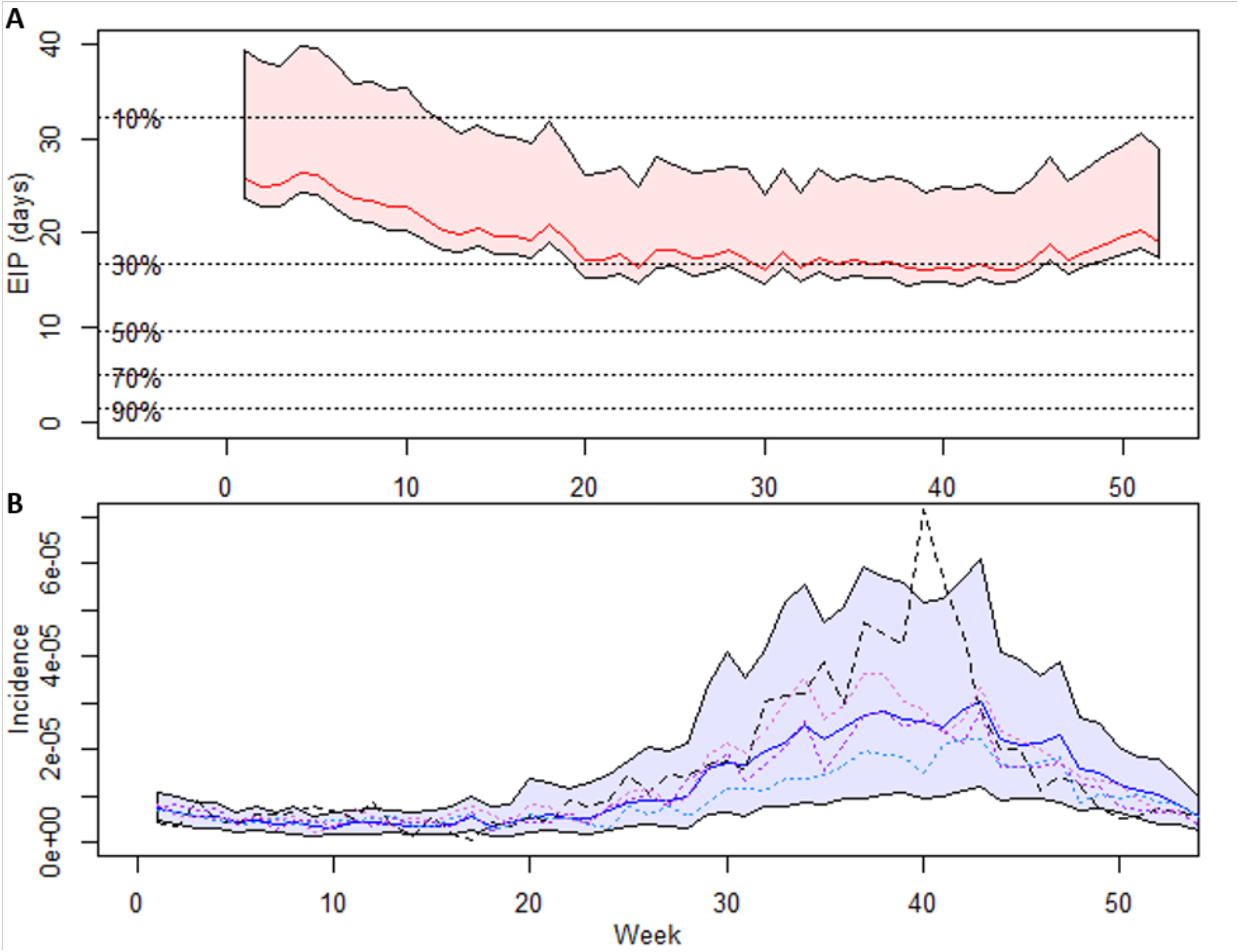
Results from an alternative calibration. Here DENV was introduced to the simulation 140 days before the simulation proper, and the starting conditions were different from that of the main article. A) The average extrinsic incubation period (EIP) at each week of the simulation, if the temperatures during that week had remained constant. The red line indicates the median across all municipalities, and the shaded region spans the minimum and maximum EIP. Dotted black lines provide the proportion of adult mosquitoes that are expected to live for at least the corresponding number of days. B) Weekly reported incidences of dengue in the San Juan Metropolitan Statistical Area between Jan 1, 2007 and Dec 31, 2007 (dashed black line) along with the median reported incidence across replicate simulations (solid blue line) and 95% confidence intervals (blue area). Colored, dotted lines are example replicate simulations.

**SFig. 4.**
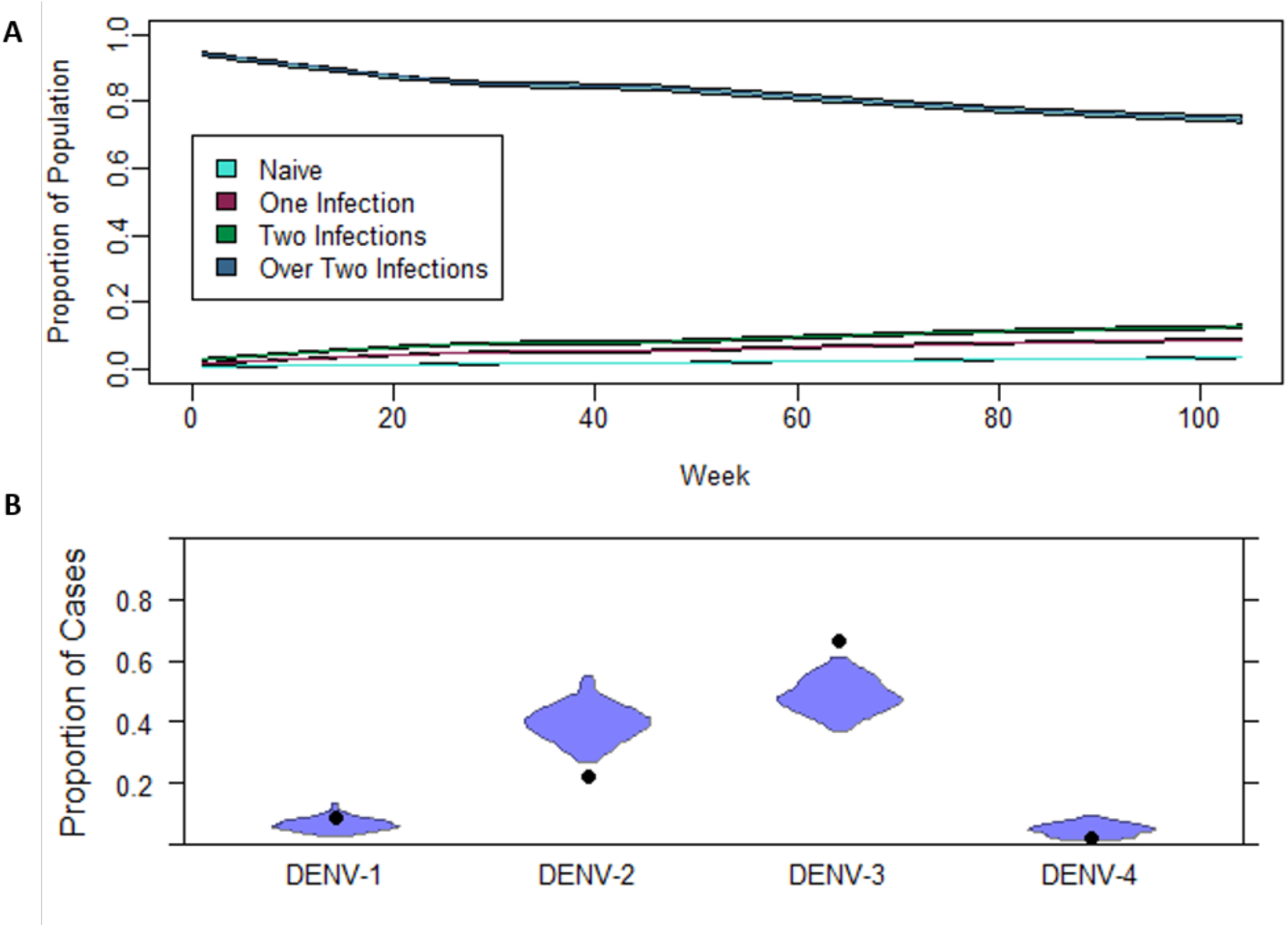
Results from an alternative calibration. Here DENV was introduced to the simulation 140 days before the simulation proper. The starting condition was the same as that of the main article. A) The median proportion of the population across replicate simulations that were immunologically naïve (cyan) or had experienced one (magenta), two (light green), or more than two (teal) previous DENV infections (solid lines), along with 95% confidence intervals (colored regions). B) Violin plots of the proportion of reported cases in a simulation attributed to each serotype. Circles indicate the proportion of cases that were attributed to each serotype in the San Juan Metropolitan Statistical Area between Jan 1, 2007 and December 31, 2007, out of those cases for which a serotype was known.

### Mosquito population dynamics

Between January 2007 and December 2008, air temperature and rainfall patterns led to fluctuations in the size of the adult mosquito population (Fig. 3A-C) and the average bites per human (Fig. 3D). The median number of daily bites per human across replicate simulations ranged from 1.053 to 2.692. Rainfall was the primary driver of local peaks in mosquito populations, with most large influxes of rain corresponding to a spike in adult mosquitoes a few weeks later. In combination with the local peaks in mosquito density associated with rainfall, the vector population also showed broader seasonal trends that were well correlated with air temperature (r = 0.638). Based on these vector dynamics and the maximum-likelihood estimate of *c_trap_* (3.65 x 10^−05^), the expected numbers of female *Ae. aegypti* caught in BG traps between Nov. 2007 and Dec. 2008 showed similar trends across months as were reported by Barrera, Amador, and MacKay (2011) (Fig. 4). Except for Feb. 2008, the expected number of mosquitoes increased and declined at the same times in the simulations as in the reference data. The magnitude of each increase or decline, however, was more mixed, with the expected number of mosquitoes in some months nearly identical to the reference (e.g., July-Sept. 2008 and Nov. 2008) and less well matched in others (e.g., Nov.-Dec. 2007).

**Fig. 3:**
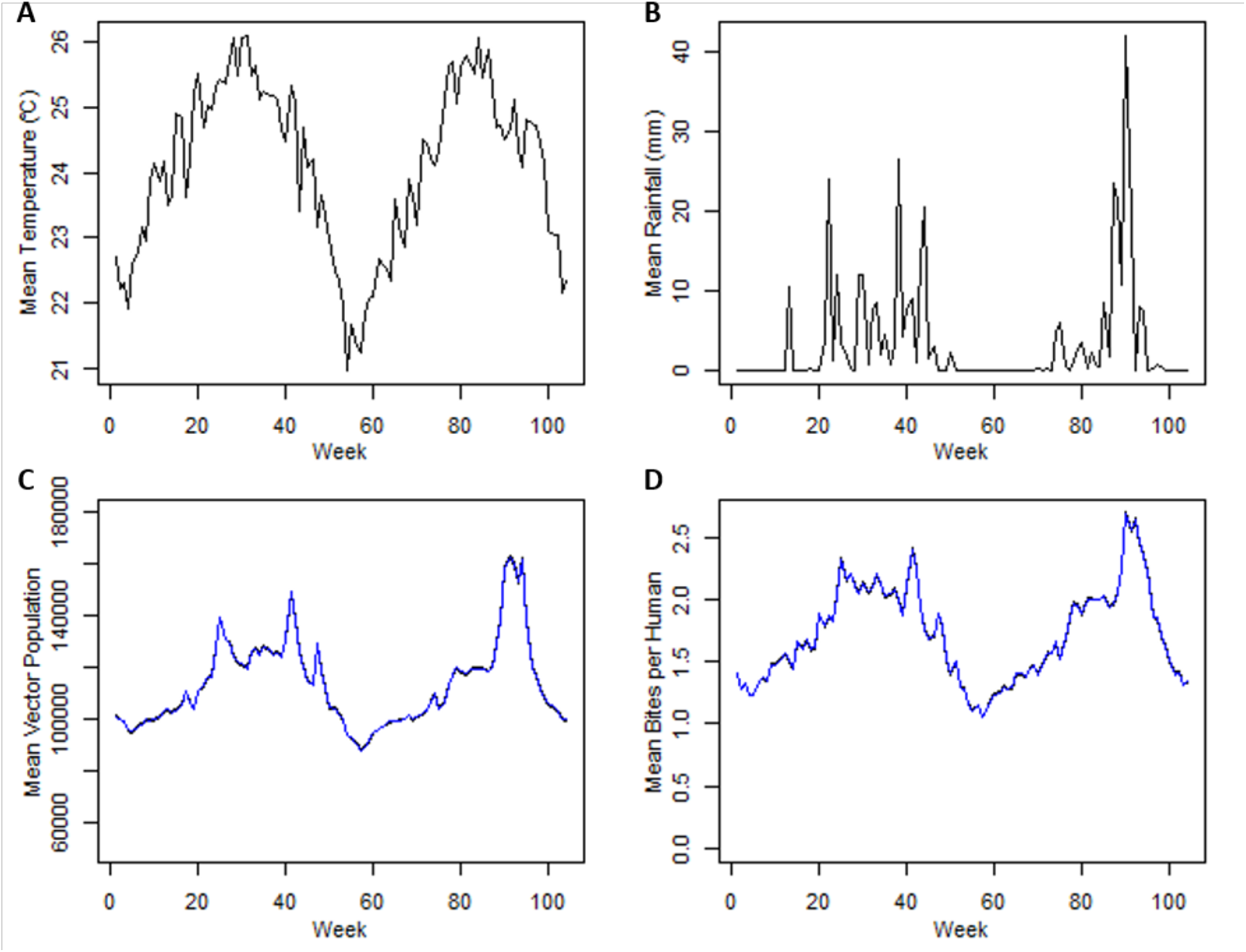
A) The weekly mean temperature and B) rainfall in the San Juan-Carolina-Caguas Metropolitan Statistical Area (MSA) of Puerto Rico between Jan 1, 2007 and Dec 31, 2007. C) The weekly mean adult vector population of the San Juan MSA based on the model. D) The weekly mean bites per human based on the model.

**Fig. 4:**
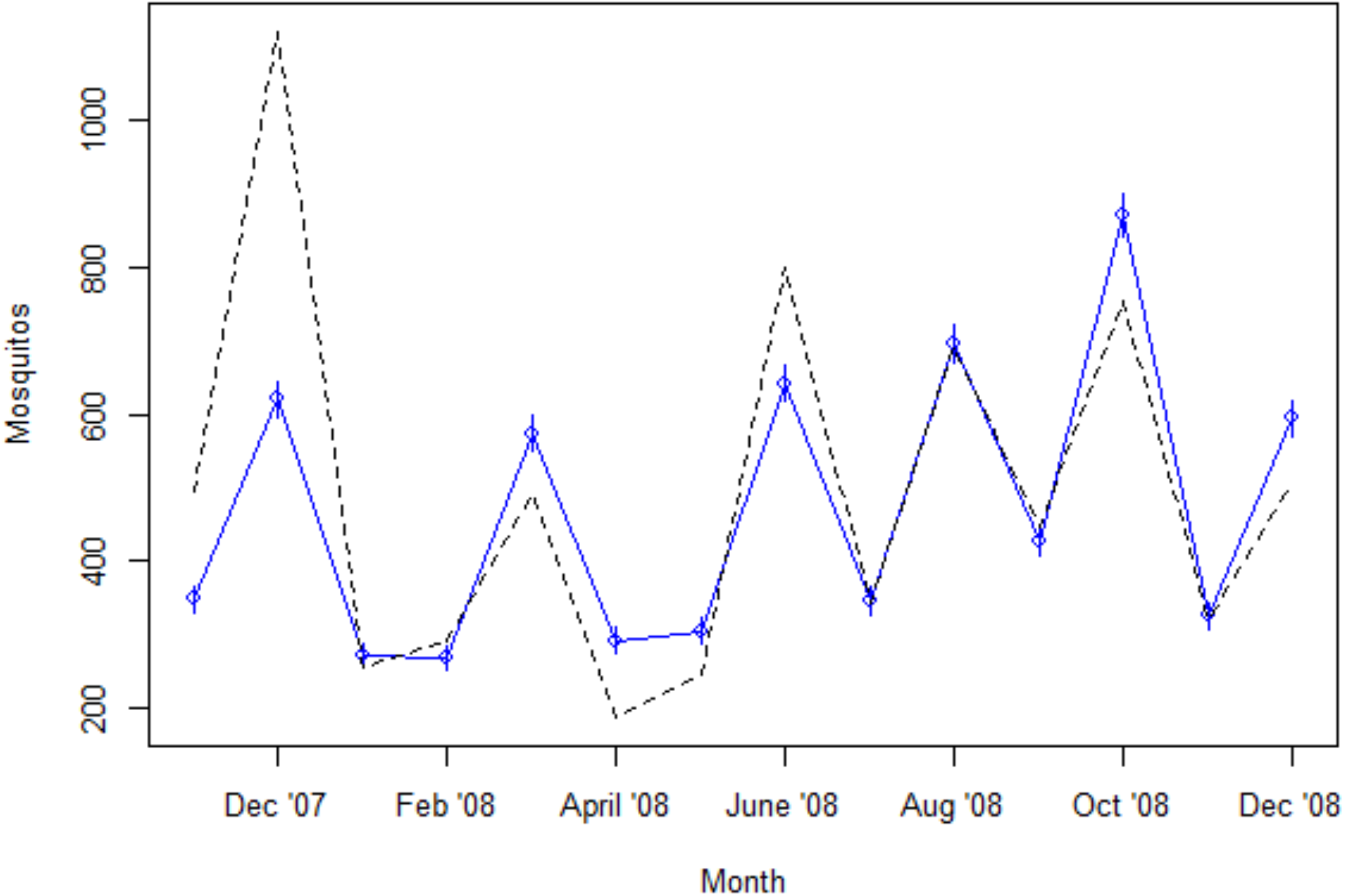
The expected number of female *Ae. aegypti* caught in BG traps between November 2007 and December 2008 based on the model parameters and the average number of adult vectors in the simulations (in blue), along with the actual number trapped in Barrera, Amador, and MacKay (2011) (black, dashed line). The number of trapped vectors is assumed to follow a Poisson distribution with an expected value equal to the product of the average number of vectors in a month, the number of traps, and a fitted scaling parameter. Bars represent one standard deviation.

The average duration of the extrinsic incubation period (EIP) varied with temperature across municipalities and time (Fig. 5). The longest EIPs consistently occurred in Orocovis, but the shortest EIPs alternated between the Fajardo and Loiza municipalities, except in the fourth week when the lowest EIP occurred in Cataño. Given the assumptions of the model, at any given time no less than 20% but no more than 60% of adult mosquitoes were expected to survive at least as long as the EIP.

**Fig. 5.**
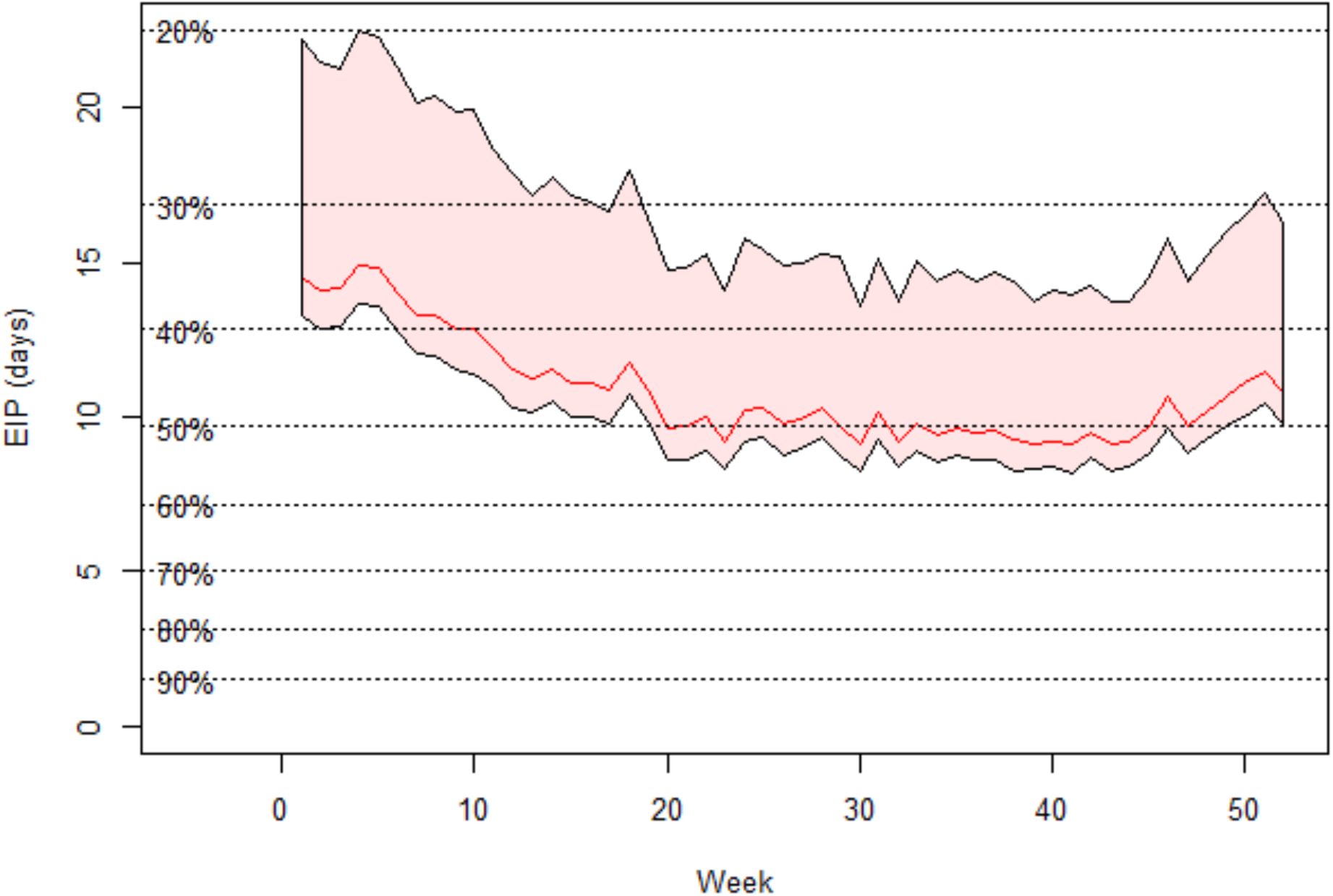
The average extrinsic incubation period (EIP) at each week of the simulation, if the temperatures during that week had remained constant. The red line indicates the median across all municipalities, and the shaded region spans the minimum and maximum EIP. Dotted black lines provide the proportion of adult mosquitoes that are expected to live for at least the corresponding number of days.

### Total incidence and infection dynamics

Reported incidence in the simulations generally corresponded well to reports from the PDSS in 2007. The 95% confidence intervals on the reported incidence included the corresponding PDSS incidence in 45 out of 52 weeks. Further, there were comparable temporal trends in incidence between the simulations and the reference data. The reported incidence of dengue in the San Juan MSA remained at a stable, low level for the first 17 weeks of 2007 before beginning an upward trend that peaked at week 40 before declining for the rest of the year (Fig. 1). Similarly, the simulations’ median reported incidences were predominantly stable and low at the beginning of the year, rose at around week 17, and had a declining trend between week 40 and the end of the year. However, unlike the PDSS data, the median incidence levelled off at around week 34 and began to decline earlier, at around week 37 (Fig. 1).

Predictions on the cumulative cases that occurred in 2007 differed between simulations (Fig. 6A). After accounting for differences between the simulated population sizes and the population size of the actual San Juan MSA, most simulations predicted between 500 and 3,000 cases with a median of 1,779 cases, although one simulation predicted as many as 3,565 cases. The median prediction was similar to the actual cumulative number of cases, 1,969 cases, leading to a relative error of 9.66%. The predicted number of infections also varied between simulations but was generally between 40,000 and 190,000 infections (median: 126,515 infections), although three predictions were over 200,000 infections (max.: 246,938 infections) (Fig. 6B). Despite the variability between predictions for the cumulative cases and infections, though, the predicted reporting rate was consistently around 1% (min.: 1.36%, max: 1.50%) (Fig. 6C).

**Fig. 6.**
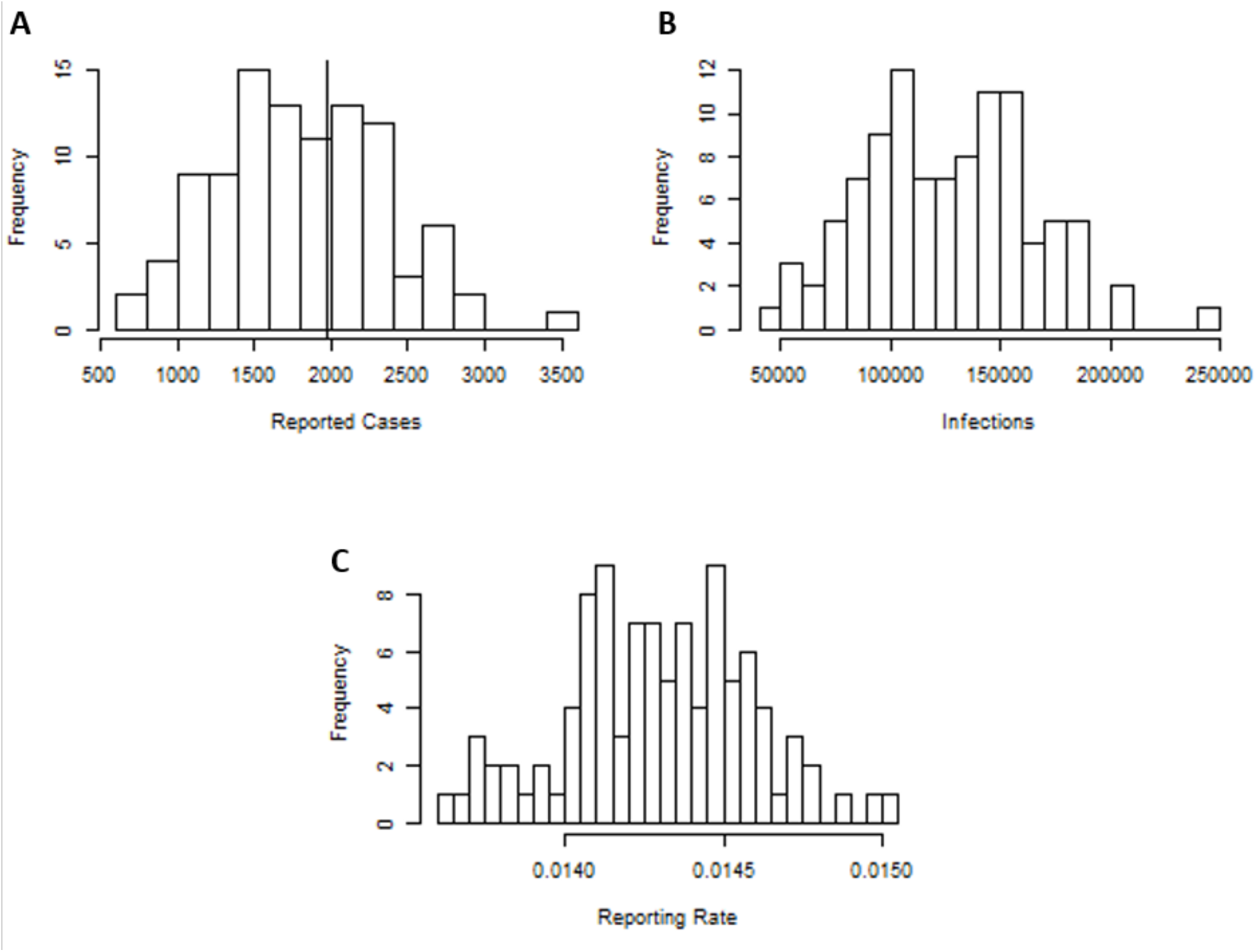
Predictions for the number of reported cases, number of infections, and reporting rate in the San Juan-Carolina-Caguas Metropolitan Statistical Area (MSA) in 2007. Each prediction is the corresponding output from one replicate simulation times a correction factor to account for the difference in population sizes between the simulations and the actual San Juan MSA. A) Reported cases, B) Number of infections, C) Reporting rate.

The disease dynamics of individual municipalities were variable but generally fell into three categories (Fig. 7). In municipalities such as Florida and Orocovis, there was very little sustained transmission (Fig. 7A,B). The median number of new cases in a week was never greater than zero, and there were no new infections a few weeks into the year. For example, Orocovis had no new infections after the ninth week. In contrast, the median number of new infections at each week in municipalities such as Caguas and Loiza faded to zero early in the year but in some simulations had continual transmission for nearly all of 2007 (Fig. 7C,D). Finally, the median number of new cases in municipalities such as Bayamon, San Juan, and Carolina follow a prototypical epidemic trajectory characterized by a distinct peak (Fig. 7E,F).

**Fig. 7.**
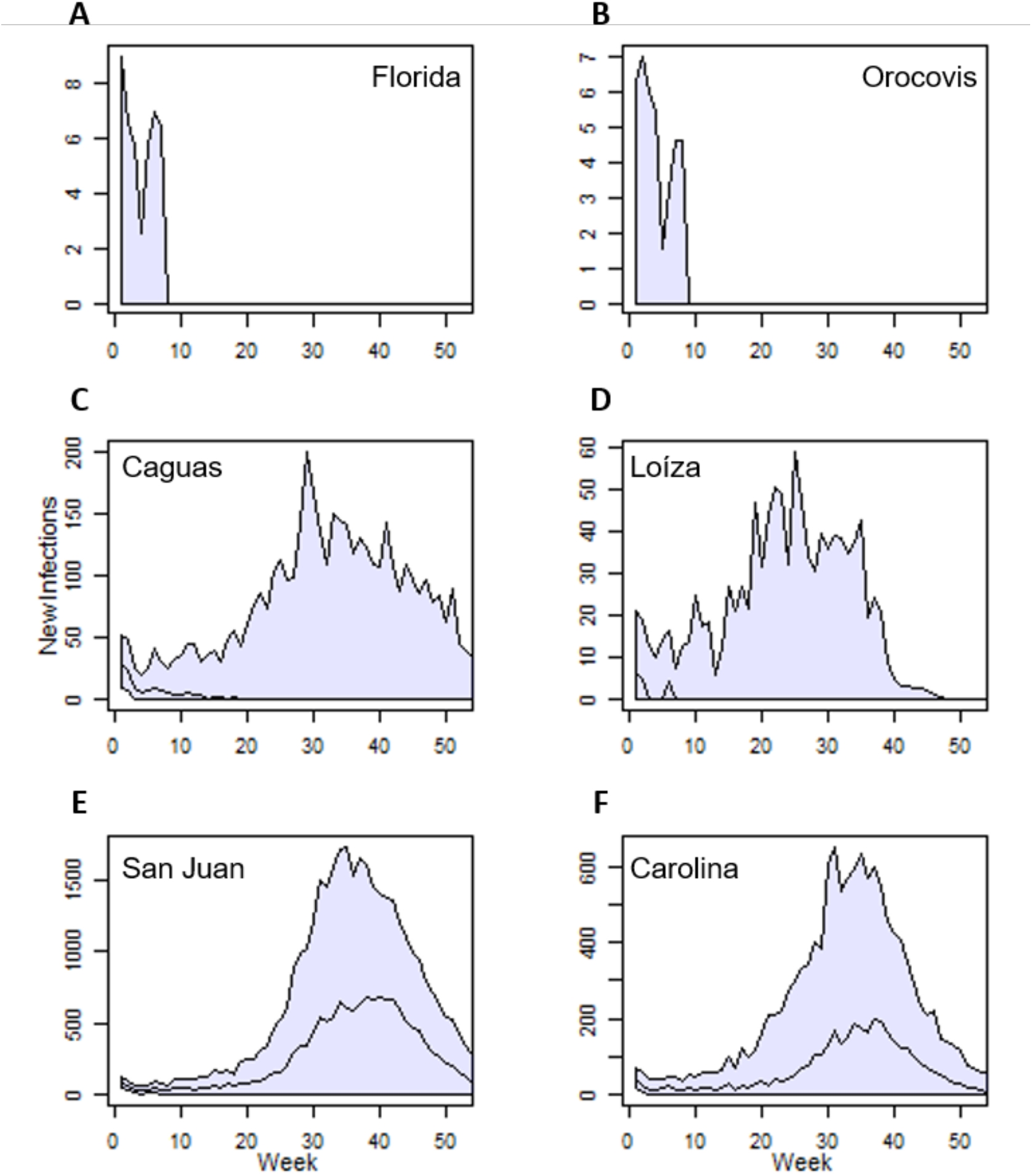
The median number of new infections across replicate simulations (solid line), along with 95% confidence intervals (blue area), for six municipalities in the San Juan-Carolina-Caguas Metropolitan Statistical Area. Plot labels provide which municipality is represented. These six exemplify the three general DENV dynamics observed in the simulations: A,B) early fadeouts in nearly all simulations, C,D) early fadeout in at least half of all simulations but with year-long transmission in some simulations, E,F) year-long transmission in over half of all simulations and with a prototypical epidemic trajectory.

### Age distribution of cases and immunity

Tomashek *et al.* (2009) reported that the age group with the most cases was 10-19-year-olds and that the share of cases declined with each subsequent age group (Fig. 2A). The 0-9-year-olds composed 10.48% of all cases, which was intermediate between 30-39-year-olds and 40-49-year-olds. For the most part, this pattern was present in both the (marginal) median and medoid predictions on the age distribution of cases, except that 10-19-year-olds and 20-29-year-olds were switched in the median and 0-9-year-olds and 40-49-year-olds were switched in the medoid (Fig. 2A). This pattern deviated from the age distribution of the population overall, which had a more even distribution across age groups and a larger representation of individuals over 70 years old (Fig. 2B). However, the exact proportion of cases associated with each age group showed less concurrence. The proportion of cases associated with 0-9-years-olds and 60-69-year-olds in the PDSS data were nearly identical to these age groups’ median in the simulations, and the proportion associated with individuals over 70 years old was just outside the 95% confidence intervals of the simulations. However, there were many fewer cases associated with 10-19-year-olds in the simulations and more cases associated with the remaining age groups.

Age and immunity were closely coupled in the simulations, with older age groups tending to be immune to more DENV serotypes than younger ones (Fig. 8). For example, at the beginning of the simulation 69.17% of all 0-9-year-olds had no immunity to any serotype. With each increasing age group, a larger proportion of the population was immune to one or more serotypes, and after the 30-39-year-olds over half of each population was immune to three or more serotypes than any fewer number. In the eldest age group, around 78% of the population was immune to three or more serotypes, and less than 1% was immunologically naïve. This pattern persisted to the end of the simulation, although there was more uncertainty about the exact proportions in each immune status (SFig. 5)

**Fig. 8.**
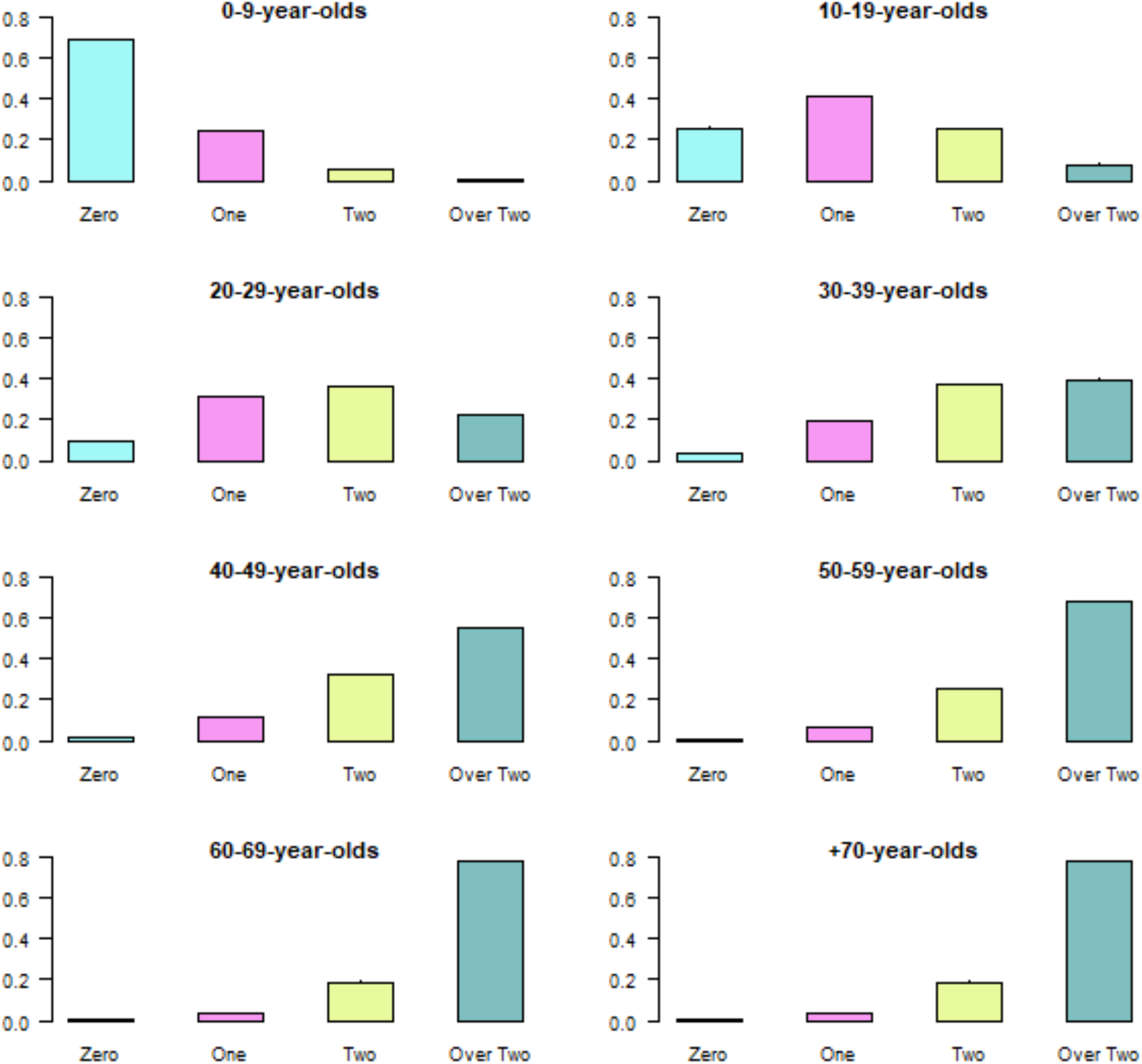
The median proportions of each age group that were immune to zero, one, two, or more than two DENV serotypes on the first day of the simulation, excluding the burn-in period. There was generally too little variability to visualize 95% confidence intervals.

**SFig 5.**
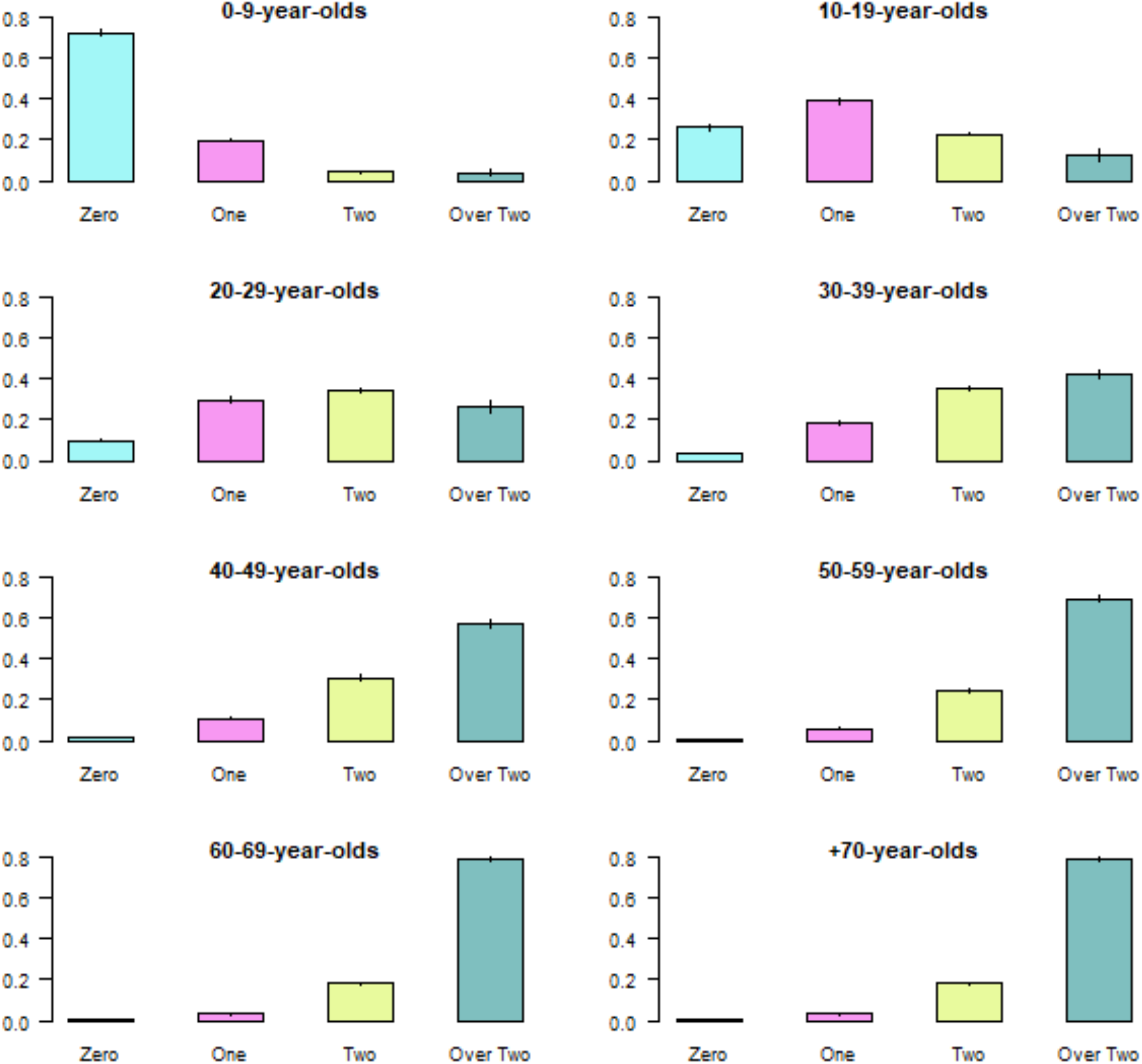
The median proportions of each age group that were immune to zero, one, two, or more than two DENV serotypes on the final day of the simulation. Bars indicate 95% confidence intervals.

The relative prominence of immune groups was stable over time (Fig. 9). The greatest proportion of the population was immune to three or more DENV serotypes, and the naïve individuals composed the smallest proportion of the population (Fig. 9A). Within these broader immune statuses, the populations had the greatest immunity to DENV-2 and the least immunity to DENV-3 (Fig. 9B). The level immunity to DENV-1 and DENV-4 were nearly identical. Although the relative prominence of immune statuses was stable, their exact proportions changed as the outbreak progressed. The proportion of the population immune to three or more serotypes grew the most dramatically because recovered individuals experienced a period of heterologous cross-immunity, regardless of their previous immune status. Since the simulation was too short to observe the cessation of cross-immunity, the median proportion of the population immune to one or two serotypes declined slightly over the course of the simulation. The naïve population remained stable, implying that the infection rate in naïve individuals was comparable to the population’s birth rate.

**Fig. 9:**
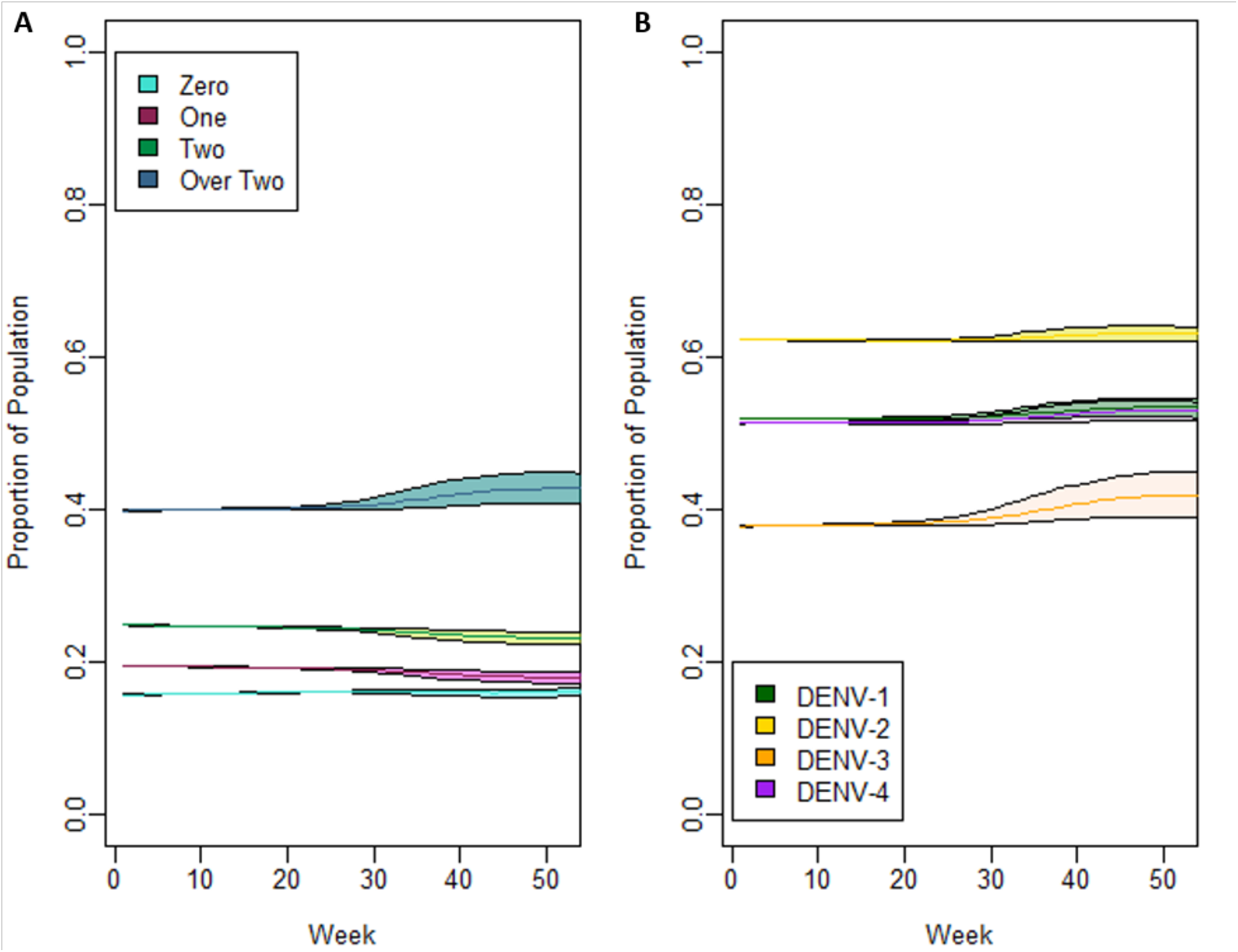
A) The median proportion of the population across replicate simulations that were immune to zero (cyan), one (magenta), two (light green), or more than two (teal) DENV serotypes (solid lines), along with 95% confidence intervals (colored regions). B) The median proportion of the population immune to each DENV serotype across replicates of the maximum likelihood particle (solid lines), along with 95% confidence intervals (colored regions). DENV-1: Green, DENV-2: Yellow, DENV-3: Orange, DENV-4: Purple. previous infections (solid line).

### Serotype dynamics

The correspondence between serotype dominance in the simulations and the PDSS data was mixed (Fig. 10). In accordance with the PDSS data, DENV-3 was generally more prominent than any other serotype, and the proportion of cases associated with DENV-1 and −4 that PDSS reported fell within the range present in the simulations. However, DENV-2 circulated at a lower proportion in the simulations than was reported in the PDSS. This discrepancy may have arisen because the simulations tended to feature dominant circulation by one serotype as opposed to the somewhat more even mixture of serotypes observed in the PDSS data.

**Fig. 10:**
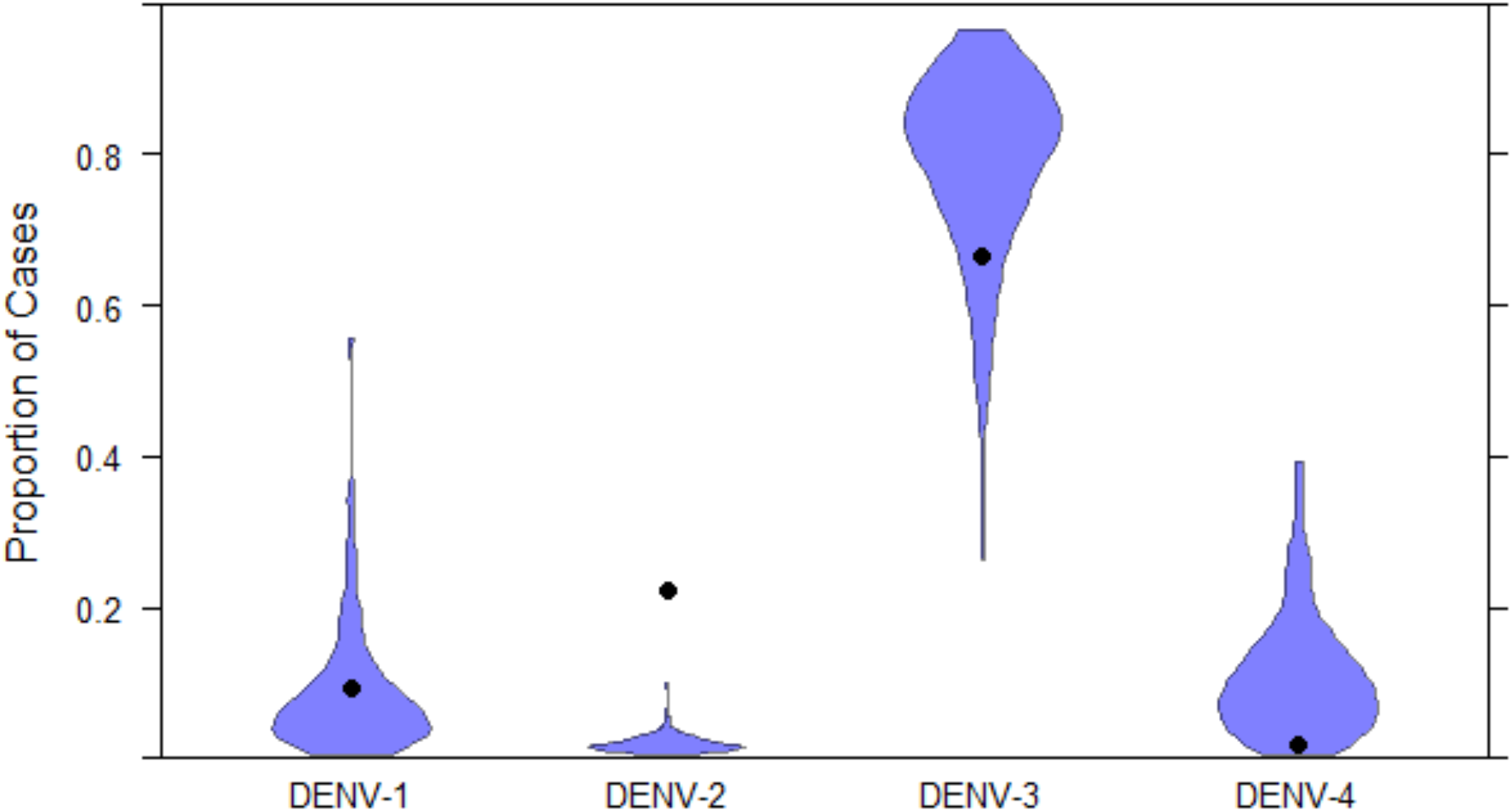
Violin plots of the proportion of reported cases in a simulation attributed to each serotype. Circles indicate the proportion of cases that were attributed to each serotype in the San Juan Metropolitan Statistical Area between Jan 1, 2007 and December 31, 2007, out of those cases for which a serotype was known.

## DISCUSSION

We have presented a new climate-driven, agent-based model for DENV transmission that incorporates detailed models of both the vector *Ae. aegypti*—including egg, larval, immature, and adult life stages—and the human host—including infectiousness, immunity, disease status, and demography. We also performed an example implementation of this model based on the San Juan-Carolina-Caguas Metropolitan Statistical Area (MSA) of Puerto Rico between January and December 2007. The model captured broad trends in multiple types of reference data, including the incidence time series, age distribution of cases, serotype distribution of cases, and vector dynamics. Not all of these data sets are required for model calibration, but our experience here suggests that they each add value to the model calibration process in their own way. Other aspects of the model’s specification draw on empirical estimates from the literature, and still others benefit substantially from adaptation of the structure of related models for other diseases (Bershteyn et al., 2018).

One aspect of the model in which the availability of empirical data for calibration was particularly useful was vector population dynamics. Barrera *et al.* (2011) reported clear interannual patterns in the population of *Ae. aegypti* that were closely correlated with air temperature and precipitation. Similarly, the population of adult vectors in our simulations were tightly connected to air temperature and rainfall inputs. On a broad scale, the number of vectors followed the seasonal dynamics of air temperature, whereas influxes of rainwater led to localized peaks in the vector population. For instance, there was an anomalously high level of precipitation at week 38 of 2008. This in turn led to the simulations’ largest vector population at week 39 of 2008.

Similarly, the model also created realistic, broad patterns in the transmission dynamics that were not directly apparent in the reference data. Municipalities with larger populations would be expected to have greater DENV transmission because their size would make them less susceptible to stochastic fadeouts, provided that a sufficient proportion of the population were susceptible. Similarly, municipalities with greater temperatures are expected to have greater transmission, in part because the EIP will be lower in these regions and therefore more mosquitoes will survive long enough to become infectious. These expected patterns are seen in the simulation results. San Juan, Carolina, and Bayamon municipalities are three of the most populous municipalities in the San Juan MSA, and all three experience relatively high temperatures. As a result, in over half of all simulations, these municipalities had sustained transmission for the entire year. When predicting disease dynamics for a municipality, the model weighs these two factors. For example, Loiza municipality had some of the highest EIPs in the simulations, yet, likely because of its smaller population, disease transmission in this municipality was more modest. A similar result occurred in Caguas, which has a large population but a cooler climate only permitted modest transmission. Still other factors not incorporated into the model could also play an important role in determining spatial patterns of dengue incidence and immunity to DENV. For example, socioeconomic conditions have been observed to correlate with transmission via their associations with factors such as the availability of breeding habitats for mosquitoes and mosquito-human contact as affected by housing characteristics (Reiter *et al.* 2003; Farinelli *et al.* 2018).

A final broad process that the model replicated well was the relationship between immunity and serotype dominance. The prominence of a serotype in a population is expected to be greater when the population has little immunity to it than when it has more. In every simulation, there was less immunity to DENV-3 than any other serotype, and this was associated with greater circulation of DENV-3 relative to other serotypes. Similarly, the population had a comparable level of immunity to DENV-1 and DENV-4 across simulations, which was associated with those serotypes circulating at similar levels. Finally, the population had the greatest level of immunity to DENV-2, and this serotype usually circulated at a lower proportion than the others.

Whereas the model performed well in extracting broad patterns, the model’s ability to provide exact predictions for epidemiological parameters was more mixed. For most weeks, the incidence reported by PDSS fell within the 95% confidence intervals established by the model, and the predicted proportion of cases associated with 0-9-year-olds, 60-69-year-olds, and individuals over 70 years old were comparable between the simulations and PDSS. However, other parameters, like the proportion of cases associated with 10-19-year-olds or with DENV-2, were not as well matched between the reference and simulated data. Constraints that are necessary to allow DTK-Dengue to easily transition between locations and times may contribute to some of these difficulties. For example, DTK-Dengue assumes that the historical force of infection is constant across all years. Because of this assumption, the user does not need to provide details about a location’s immunological history and therefore can transition between locations with greater ease. However, this assumption strongly constrains the relationship between age and seropositivity, which makes the age distribution of cases more difficult to replicate. For example, the San Juan MSA had a major dengue outbreak in 1994 whose cumulative incidence was 43% greater than any other outbreak between that year and 2007. This outbreak would have likely had a major influence on the immune status of individuals over the age of 13 in 2007 but would have been absent in those younger than 13. Yet, since DTK-Dengue assumes that the historical force of infection is constant, our model would have had to strike a balance between these two groups. Under this consideration, it is somewhat logical that the model under predicts the incidence of dengue in 10-19-year-olds but over predicts its incidence in 20-59-year-olds. In future implementations, the model may be better able to replicate the age distribution of cases if it used a temporally-variable force of infection, similar to Quan *et al.* (2018).

In addition to the constraints that allow DTK-Dengue to switch between settings easily, the breadth of topics about which it provides information can also limit the precision of its predictions. With such a large variety of outputs, the model must strike a compromise between realism amongst each one. In some cases, the model has been able to provide better constrained predictions for some output streams if other output streams were less realistic. For example, when the extrinsic incubation period was allowed to be longer, the model more accurately predicted the peak week in incidence. Similarly, when immunity was allowed to reach extreme levels, the dominance of each serotype became better constrained.

Finally, some loss of precision likely arises from imperfect knowledge about the system. This could have particularly impacted the serotype-specific incidences in the simulations. Although the model identified that DENV-3 was the most prominent serotype, its dominance was generally overestimated and frequently nearly all cases were associated with it. The discrepancy between model results and the PDSS data could originate in differences in serotype infectiousness or in the probability that infections become symptomatic. The degree of these differences, if they exist, is not well established, so they are currently difficult to incorporate into the model.

Although the breadth of topics discussed here is highly diverse, EMOD-DTK, and by extension DTK-Dengue, can incorporate a great deal more features as well. One particularly promising feature are campaign events. These model features allow a researcher to explore how proposed DENV and vector control strategies would impact an affected area. For example, there are campaigns to explore the effectiveness of indoor residual spraying and increased barriers to entering dwellings. These tools can assist policymakers in deciding where to invest into control campaigns to maximize benefit and the expected effect size of each decision. With these applications in mind, DTK-Dengue is a promising new tool in the study of arboviral diseases.

